# Programming super DNA-enzyme molecules for on-demand enzyme activity modulation

**DOI:** 10.1101/2022.09.26.509444

**Authors:** Haipei Zhao, Xuehao Xiu, Mingqiang Li, Mingyang Gou, Leyang Tao, Xiaolei Zuo, Chunhai Fan, Zhongqun Tian, Ping Song

## Abstract

Dynamic interactions of enzymes, including programmable configuration and cycling of enzymes, play important roles in the regulation of cellular metabolism. Here, we construct a super DNA-enzymes molecule (SDEM) that comprises at least two cascade enzymes and linked DNA strands to control and detect metabolism. We find that the programmable SDEM which comprises glucose oxidase (GOx) and horseradish peroxidase (HRP) has a 50-fold lower detection of limit and a 1.6-fold higher reaction rate than free enzymes. SDEM can be assembled and disassembled using a hairpin structure and a displacement DNA strand to complete multiple cycles. An entropically driven catalytic assembly (catassembly) enables different SDEMs to switch from SDEM with GOx and HRP cascades to SDEM with sarcosine oxidase (SOX) and HRP cascades by over six orders of magnitude less time than no catassembly to detect different metabolisms (glucose and sarcosine) on demand.

## Introduction

Metabolon was defined as a supramolecular complex of sequential metabolic enzymes and cellular structural elements by Paul Srere in 1987^1^. The spatial and temporal organization of enzymes in metabolons, i.e. multi-enzyme complexes participating in cascades reactions, is important to regulate the metabolism^2^. Hence, the ability of fine-tuned metabolons is crucial to maximize the production of metabolism. Inspired by nature, artificial protein and DNA scaffolds have been used to co-localize enzymes in a controlled manner to enhance metabolic reactions^3-7^. For example, with the advent of artificial proteins and DNA origami, it became possible to colocalize multiple enzymes efficiently^3^, and enable enzymatic co-assembly precisely^8,9^. Moreover, DNA nanocages have been used to encapsulate various enzymes in nano-confined cavities to reduce the interference of proteases, and improve catalytic activity as well as stability ^10^. The above methods have significant advantages in the co-assembly of enzymes. However, those methods still cannot achieve dynamic regulation.

Dynamic regulation of enzymes is the response from the life to external stimuli in nature^11,12^. For example, purinosomes are mesoscale assemblies and have dynamic and reversible structures which can face the purine depletion in mammalian cells^2^. In addition, phosphorylation of the purinosomes by the CK_2_ kinase promotes its dissociation, whereas assembly can also be initiated by phosphatases or kinase inhibitors^2,13-15^. Dynamic regulation of enzymes in response to the external environment enables cells to maintain cellular activity^14^. Similarly, dysregulation of enzymes also promotes cancer development and progression^16,17^. In vivo, enzymes are stable or transient complexes function with other enzymes^18-20^ so that enzyme regulation plays a key role in health. Programmable DNA nanotechnologies have been used in enzyme dynamic regulation based on toehold-mediated strand displacement^21-27^. Compared with genetic circuit designs limited by the degradation of the associated biological components such as proteins and RNAs^28-30^, the regulation of enzymes by toehold-mediated strand displacement is much faster. For example, DNA tweezers were used to control dual-enzyme cascades efficiently by regulating the switchable distance between open and closed conformations^31,32^. Recently, the regulation of proteins by toehold-mediated strand displacement achieved not only the regulation of proteins by complex and diverse logic gates, but also achieved the dynamic regulation of enzyme activities in vivo^21,22^. However, this is only within one metabolon of dual-enzyme assembly and disassembly and is sensitive to only one metabolism. In vivo complex environment, the metabolon may have more than two enzymes which can be sensitive to several metabolites.

Here, super DNA-enzymes molecule (SDEM) was constructed to sense rare analytes highly responsively with recyclable and programmable reaction. Those structures enabled rapid and dynamic regulation of enzymes catalytic activity. Compared with the output of the fluorescence intensity of proteins, SDEMs achieved dynamic regulation among nucleic acids, enzymes, and metabolites. The reaction rate is ten times faster than the rate in the literature using a five times lower concentration of enzyme^21^. We also used toehold mediated strand displacement to conduct catalytic assembly from one set of SDEMs to a new set of SDEMs to detect different metabolism on demand.

## Results

### Programmable super DNA-enzymes molecule (SDEM) construction

We first synthesized SDEMs, which contains at least two cascade enzymes and three linked single-stranded DNA (ssDNA) fragments. Here we demonstrate using an example with glucose oxidase (GOx), horseradish peroxidase (HRP), and three ssDNA fragments. The cartoon structures of GOx and HRP are shown in Fig. 1a. The molecular weights and sizing information of GOx, HRP, and sarcosine oxidase (SOX) are shown in Figure S1-1. The SDEM(GOx-HRP) is a molecule which comprises of ssDNA (A B) binding GOx and HRP and a complementary DNA strand C (Fig. 1b). To evaluate the enzyme cascade activity, we used 3,3’,5,5’-Tetramethylbenzidine (TMB) as a medium. The SDEM(GOx-HRP) can catalyze glucose (GO) and O_2_ to produce gluconolactone and H_2_O_2_. Since the production of H_2_O_2_ is the substrate of the HRP, the cascade reaction transforms the TMB from reduced form (light blue) to oxidized form (dark blue).

**Figure 1.**
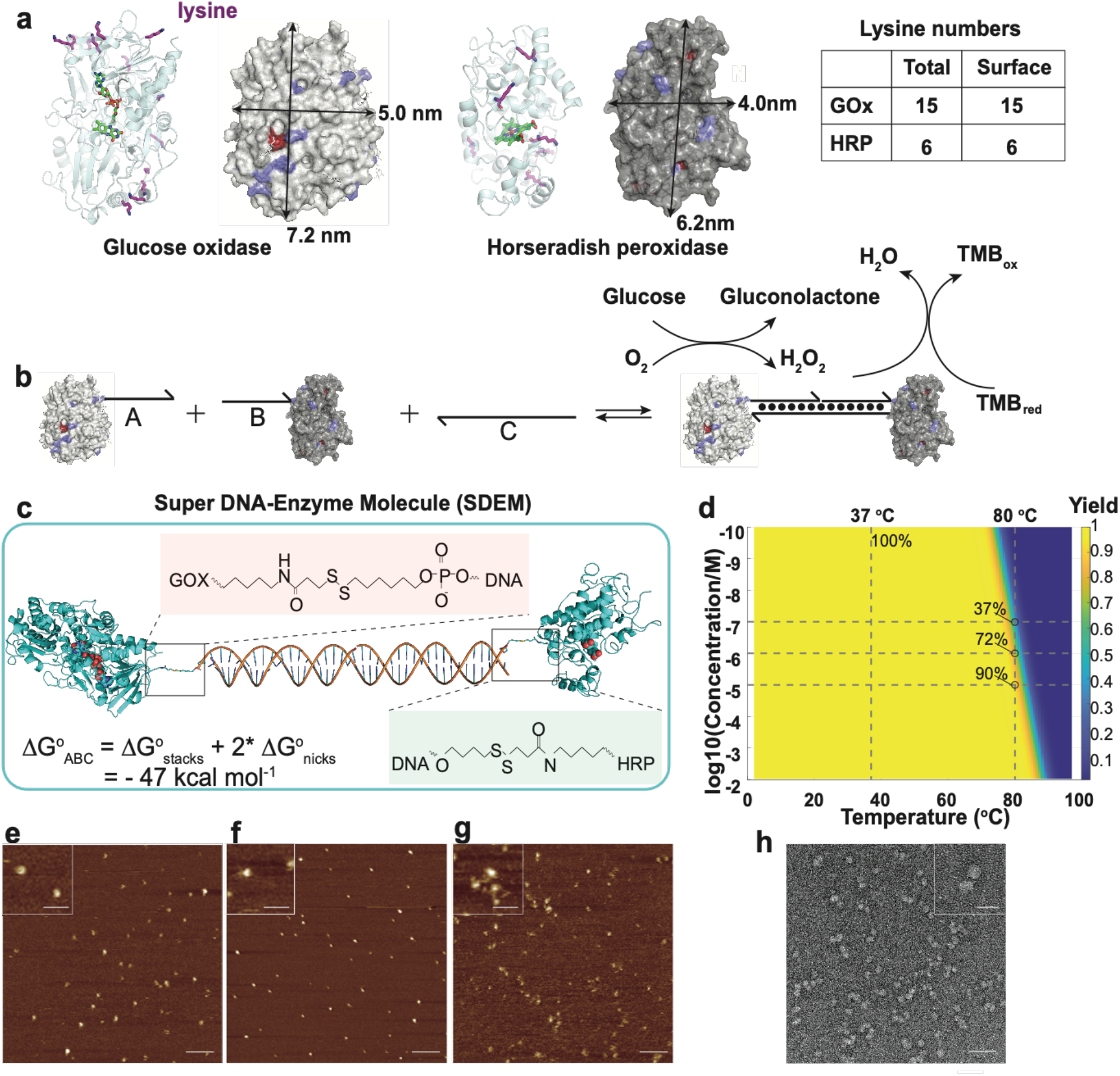
Construction of programmable super DNA-enzymes molecule. a. Cartoon structures of glucose oxidase (GOx) and horseradish peroxidase (HRP). All the lysine residues are shown as purple sticks. The activity centers for both enzymes are shown as green molecule sticks. The sizes of two enzymes are shown with spherical surface. b. Enzyme modified with DNA strands, single strand A and B can hybridize with C to form a super DNA enzymes molecule. This molecule can catalyze glucose to gluconolactone and hydrogen peroxide(H_2_O_2_), H_2_O_2_ can be catalyst by HRP and cause the TMB reduced with color change. c. The structure of GOx-DNA-HRP super molecule. The chemical structures of the linkers between enzymes and DNA are shown in the pink box and green box. The standard Gibbs energy of the double-stranded DNA is – 47 kcal mol^-1^ at 37 °C. d. The yields of SDEMs at different temperatures and different concentrations. e-g are the AFM pictures of free GOx and HRP, SSDMEs(GOx, HRP), and SDEM(GOx-HRP). The scale bar is 100 nm and insert pictures’ scale bar is 50nm. h. Negative stained TEM of SDEM(GOx-HRP). Scale bar is 50 nm and insert figure’s bar is 20 nm.

We describe the construction detail of SDEM(GOX-HRP) in Figure 1c. The lysine is an alpha-amino acid that is used in the biosynthesis of proteins^33,34^. It contains an alpha-amino group, and an alpha-carboxylic acid group, and a side chain lysyl. Wherein side chain lysyl is easy to conjugate with thiolmodified DNA (SH-DNA)^35,36^ (Fig. S1-1 to Fig. S1-3). The ratios of linked SH-DNA and enzymes are measured by ultraviolet-visible spectroscopy (Fig. S1-4). We summarized all the lysines that resides in two enzymes including on the surface numbers and total numbers in Table S1-1. All the lysines are on the surface of both enzymes. Flavin adenine dinucleotide (FAD) is the activity center of GOx in the center of the enzyme. We indicated the FAD with sticks using Pymol software (Fig. 1a). Heme activity center is also in the center of the HRP and lysine residues are all on the surface of the HRP. Thus, the modification of lysine residues will cause less activity loss since all the lysine residues are on the surface of the enzymes^8,37^.

To demonstrate the SDEM is a molecule, we calculated the binding standard Gibbs free energy of the three ssDNA at 37 °C (Fig. 1d and Fig. S2-1, Fig. S2-2). The calculation is based on stacks and nick energy with initiation penalty^38^. We calculated the standard Gibbs free energy of −47 kcal mol^-1^ which makes the structure quite stable (Fig.S2-3 to Fig.S2-7). We next simulated the hybridization yields of the three ssDNA (Fig. 1d). The hybridization yields of different concentrations from 10^−2^ M to 10^−10^ M are all 100% at 37 °C. Not surprisingly, when the temperature went up to 80 °C, we observed that the yields which varied from 37% to 90% along with the concentration of the hybridized DNA from 0.1 µM to 10 µM, even though all the strands were the same. Since most enzymes work at 37 °C, we are able to create a programmable SDEM with 100% yield at 37 °C. Notably, the construction of SDEM(GOx-HRP) was achieved and validated by Atomic Force Microscope (AFM) and negative-stain Transmission Electron Microscopy (TEM). Compared with the distances between GOx and HRP in free enzymes (Fig. 1e) and in Single Strand DNA Modified Enzymes (SSDMEs) (Fig. 1f), the distances of SDEM(GOx-HRP) (Fig. 1g) are the smallest (Fig. S2-6). We calculated the yields of SDEM(GOx-HRP) by statistically summarizing multiple data which are about 52.2% (Fig. 1h and Fig. S2-8, Table S2-1). The yield of SDEM(SOX-HRP) is shown in supplementary section S2 (Fig. S2-9 to Fig. S2-10).

### Simulation of SDEMs activity quantification

Enzyme engineering may influence the enzyme activities^39^. Meanwhile, the distances of cascade enzymes may also influence the cascade enzymes activities^8,37^. To quantify the activities of the enzymes in SDEM, we performed both experiment and simulation considering the distances of the enzymes in cascade reaction.

We firstly simulated the distances of two enzymes of free status and SDEM using GOx and HRP cascade reaction as a model (Fig. 2a). The initial distances of free enzymes and SDEM(GOx-HRP) are the same of 22.6 nm between the centers of GOx and HRP. During the diffusion, the distances of free enzymes are generally larger than the distances of enzymes in SDEM (310 k, 15.4 mM Na^+^, 5 mM Mg^2+^). Limited to the calculation speed and computer capacity, we only simulated the movement in 100 ns running on 25 nodes with total 1000 cores for three days. At 100 ns, the distance between the two enzymes is 25.5 nm for free enzyme and 14.5 nm for SDEM(GOx-HRP). The distance difference change *Δ*D (calculated in equation 1) is about 50% of SDEMs and free enzymes in 100 ns compared with 0 s.

**Figure 2.**
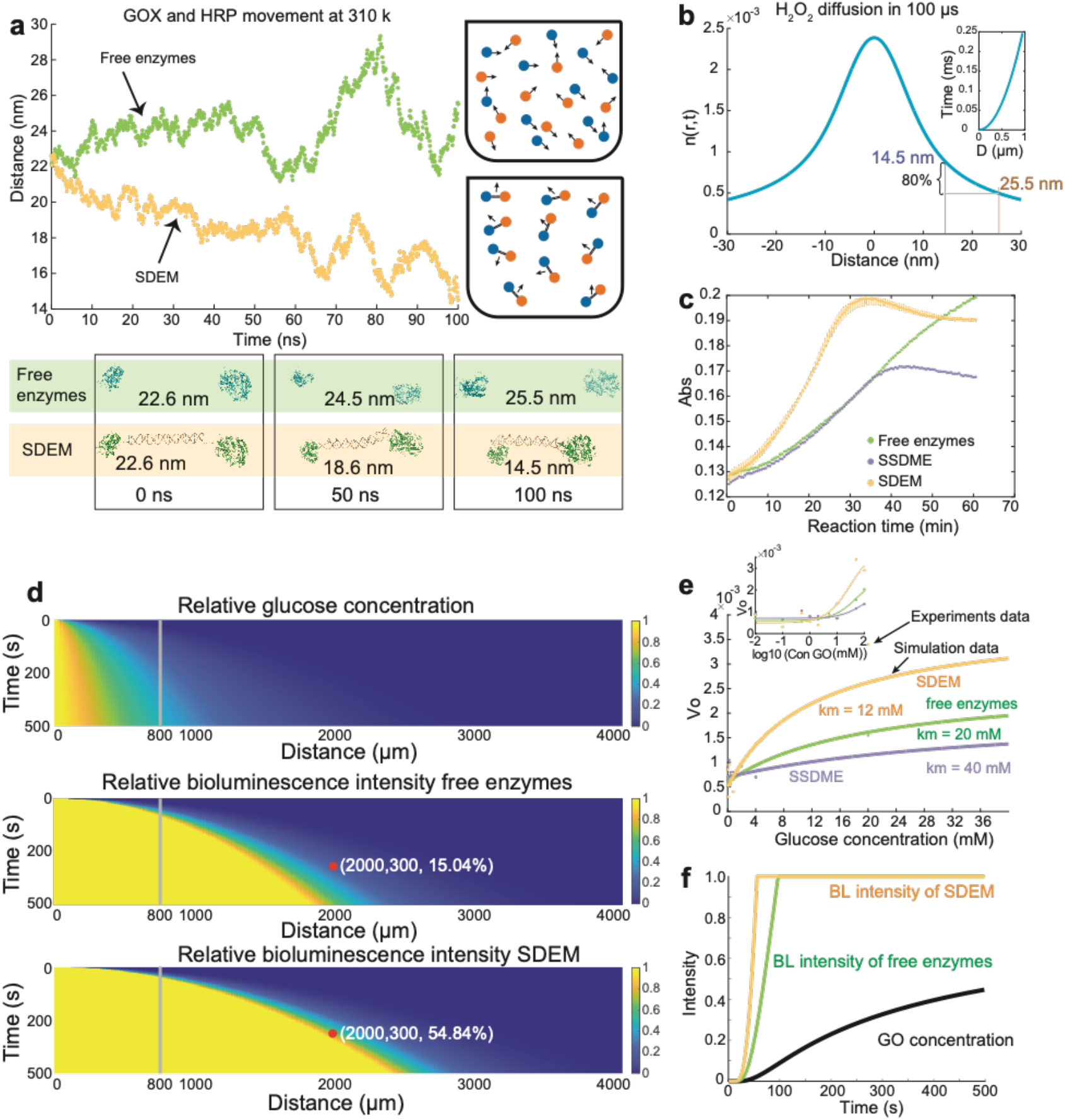
Distance and reaction activity comparison of free enzymes and SDEMs. a. MD simulation using GRACOS software. The distance between two centers of enzymes of SDEM is 22.6 nm. We also set the initial distance of two free enzymes as 22.6 nm. We simulated the distance between two enzymes of free enzymes (green dots) and SDEM (yellow dots) at 100 ns. The blue dotes and orange dotes represent the GOx and HRP in the black tubes. b. H_2_O_2_ diffusion distance in 100 µs from 3D Brownian diffusion motion, diffusion coefficient is approximately 1200 µm^2^/s at 310 K. The concentration of H_2_O_2_ when enzymes distance is 14.5 nm is 80% higher than the concentration from enzymes distance at 25.5 nm. The insert figure is the time distribution versus different distances. c. Three enzymes system detected glucose by TMB with UV absorbance at 370 nm under same concentration of GO at 4 mM and HRP at 0.8 nM, free enzyme (green dots), modified enzymes with single-stranded DNA (purple dots), SDEM (yellow dots). d. The simulation of relative glucose concentration at different time and distances using diffusion function (top). The relative bioluminescence intensity of free enzymes and SDEMs are shown in the middle and bottom using enzyme reaction function. e. The three enzyme systems reaction velocities of different glucose concentrations. The dots are the experimental data and the lines are the simulation data. The insert figure is the velocities versus log scale of different GO concentrations. All the enzymes concentration of here are 1 nM and the reaction temperature is 37 °C. f. Summary of intensity values for different time in three parameters described in d from top (black line), middle (green line), to bottom (yellow line).

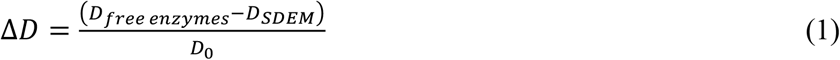

The intermediate H_2_O_2_ between enzymes of GOx/HRP cascade is essential to the cascade activity^8,37^. Here, we used Brownian motion, described by equation 2, to simulate the distance-dependent, three-dimensional (3D) diffusion of H_2_O_2_ between enzymes.

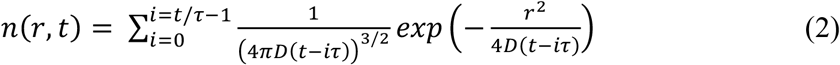

The concentration of H_2_O_2_ with distance of 14.5 nm is 80% higher than the concentration of enzymes with the distance of 25.5 nm (Fig. 2b). Not surprisingly, the difference will be changing at different simulation time. We also simulated the correlation between the diffusion distance and diffusion time in the insert figure of Figure 2b. The time scale is from 0 to 0.25 ms and distance is close to 1 µm. In the experimental detection, we usually use minutes or hours as reaction time, therefore the H_2_O_2_ diffusion time (ns to ms) won’t be a key factor to limit the final detection performance but will influence the reaction kinetics.

The reaction kinetics resulting from a different aliquot of 4 mM GO sample are shown in Figure 2c for comparison. See Supplementary section 3 for the enzymes cascade activity experiments quantified by using the absorbance of TMB at 370 nm (Fig. S3-1 to Fig. S3-4). Despite SSDMEs having similar absorbance change within 30 min as free enzymes considering the modification making a minor change of enzyme activity, the SDEMs exhibited higher absorbance change values than free enzyme in first 30 min because the distances of the two enzymes were controlled within the distances with higher H_2_O_2_.

Figure 2d displays the relative glucose concentration for diffusion time and distance. The relationship among the relative concentration of GO, diffusion distances, diffusion time can be described mathematically with diffusion function:

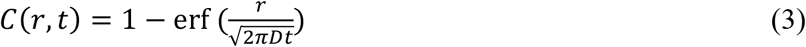

Wherein D = 6.73*10^−10^ m^2^/s, K_m_ = 20 mM, r = 0-4000 µm, t = 1-500 s.

The enzyme reaction kinetics function can be calculated with the following Michaelis-Menten equation:

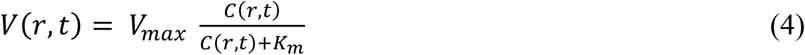

Figure 2e shows the K_m_ simulation of experiments of free enzymes, SSDMEs(GOx, HRP), and SDEM(GOx-HRP) calculated under different GO concentrations at 37 °C. The correlation between log scale of GO concentration and velocity is shown in the insert figure of Figure 2e to demonstrate the simulation of lower GO concentration.

Note that relative bioluminescence intensity of free enzymes and SDEMs are not linear. For example, the diffusion time of SDEM is shorter than free enzymes when the relative bioluminescence is close to 1 at 800 µm. To determine the correlation of intensity and time of free enzyme and SDEM system, we ran simulation on relative intensity of bioluminescence and concentration at 800 µm distance (Fig. 2f). Notably, at distance of 2000 µm and diffusion time of 300 s, the relative of bioluminescence of SDEMs is 54.84%, which is much higher than 15.04% of the free enzymes. Thus, the SDEM improves the enzymes activity compared with free enzymes because it controlled the distance of the enzymes within 22.6 nm instead of random diffusion motion.

The cascade activity of SDEMs is higher than free enzymes, with 2-fold lower K_m_ values. Since SDEM uses only three strands of DNA instead of complex DNA nanostructure, the cascade activity enhancement should mostly come from the surface-limited diffusion of H_2_O_2_ between closely spaced enzymes. Zhang et al mentioned the pH near the surface of negatively charged DNA nanostructures is a key factor to enhance the enzyme activity^40^. To test the pH effects from high concentration of DNA strands, we measured a series of higher DNA concentrations and found only very high DNA concentration like 10 µM will significantly change the enzyme activity (see Supplementary Section 4, Fig. S4-1 to Fig. S4-3). In our SDEM system, the concentration of total DNA strands is 3 nM which contributes little to enzyme activity so that the influence of DNA in SDEM can be ignored.

### Character enzyme cascades activity of SDEM(GOx-HRP)

The cascade enzymes activities were statistically studied in supplementary section S3 (Fig. S3-2 to Fig. S3-4). We next applied the free enzymes, SSDMEs(GOx, HRP), and SDEM(GOx-HRP) methods to detect GO with low concentrations (4 µM – 400 µM) in Figure 3a – Figure 3c. Due to the variation of the background signals, we were not able to detect the 4 µM GO on the three systems. The detection results of enzyme cascade activity from 4 µM to 40 mM are shown in Figure 3d (free enzymes) and Figure 3e (SDEM(GOx-HRP)). In SDEM(GOx-HRP) system, the concentration of 40 µM and 200 µM can be statistically distinguished between every two lower and higher GO concentrations. By contrast, with free enzymes and SSDMEs(GOx, HRP), the distinction between every two lower and higher GO concentration is not consistent (Fig. 3f). For example, in free enzymes group, there is a difference between 0 µM and 40 µM, 4 µM and 40 µM, but no statistical difference between 40 µM and 200 µM. In addition, there is no statistical difference between 0 µM and 200 µM so the GO concentration with 200 µM cannot be detected by free enzymes system.

**Figure 3.**
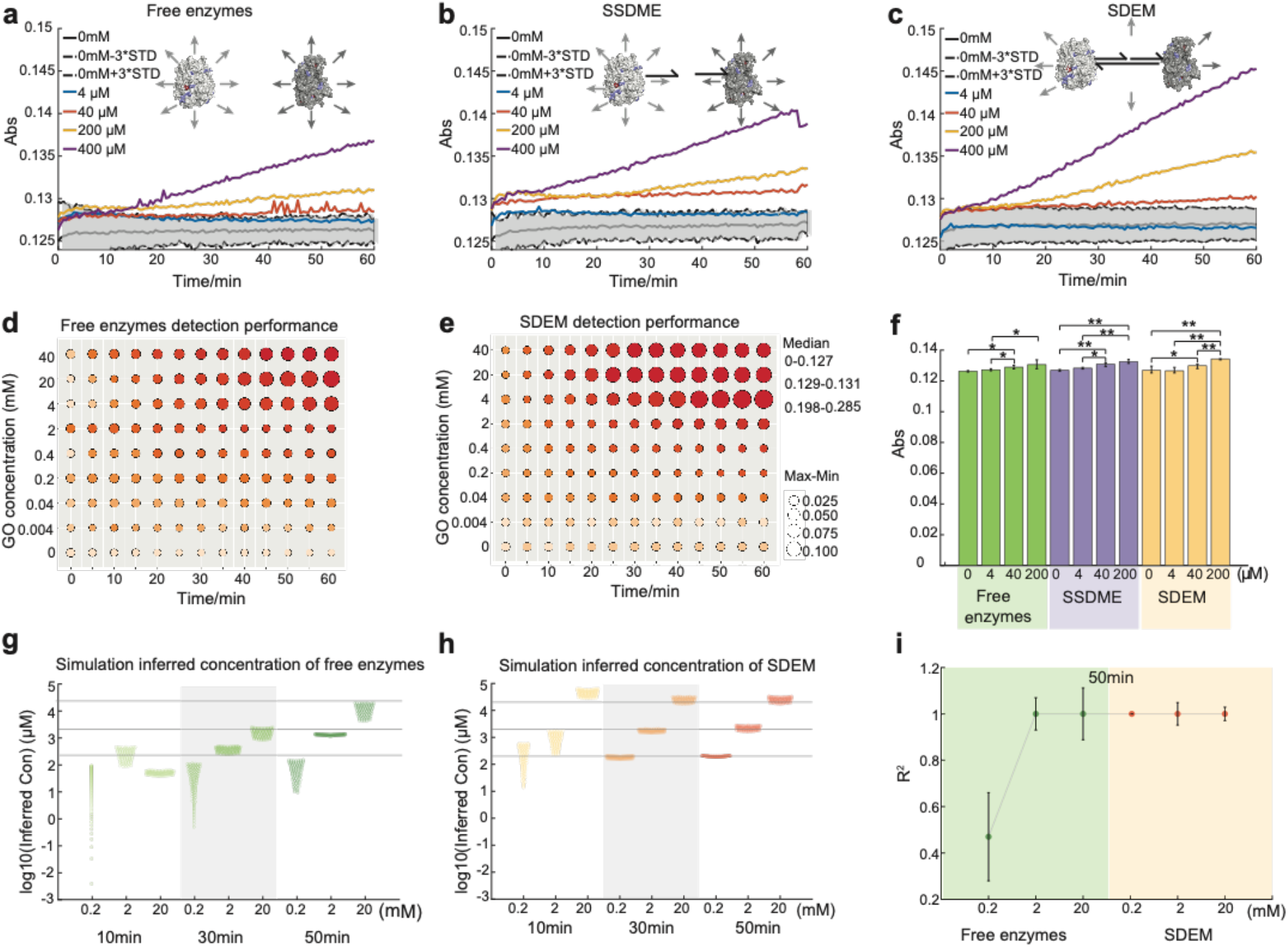
Detection and quantitation of GO concentrations based on observed UV absorbance of TMB from three enzyme systems. The lower concentration of GO (from 4 µM to 400 µM) detection used free enzymes (a), SSDMEs (b), SDEM (c). We colored the blank GO Abs and +-3 times standard deviation (STD) as gray color. GO detection using free enzymes (d) and SEDMs (e) with different reaction time. The colors of heatmap from light pink to dark red are UV absorbance values from 0 to 0.285. The size of circles represents the ΔUV absorbance values of max – min. f. Statistical analysis of three enzyme systems performance under lower GO concentration. The statistical differences between free enzymes and SSDMEs are not consistent while the SDME is consistent. Simulation of 100 inferred concentrations of GO at 0.2 mM, 2 mM and 20 mM (from lighter color to darker color) using quantification functions in reaction time of 10 min, 30 min, and 50 min of free enzymes (g) and SDEM (h). The lines from bottom to top refer to GO concentrations of 0.2 mM, 2 mM and 20 mM. The gray box highlights the inferred GO concentration at 30 min. i. Analysis of the accurate quantitation of GO using free enzymes and SDEM. Each point shows the mean coefficient of determination (R^2^) values for the inferred concentration of GO versus the real concentration at reaction of 50 min with free enzymes (green box) and SDEM (yellow box) (n=100 for each sample size). The error bars show one standard deviation. At 0.2 mM GO concentration of SDEM system, the R^2^ value reliably converged to above 0.99.

To explore the limit of detection (LOD) for free enzymes and SDEM(GOx-HRP), we increased the GO concentration up to 40 mM at different time. The free enzymes can detect 2 mM GO with relatively lower max-min values than 0.4 mM GO concentration (Fig. 3d-e). While the SDEM(GOx-HRP) has a LOD at 200 µM (Fig. 3f) which is 50-fold lower than the LOD of free enzymes. Notably, the limits of detection are related to reaction time since the colors are changed with the reaction time and the errors (max-min) are also related to the reaction time. We summarized the detection performance with different reactions and fitness functions based on the standard curves (Supplementary Section 5 (Fig. S5-1 to Fig. S5-3)).

We calculated the inferred concentration in different reactions (Fig. 3g and 3h). In 30 min reaction, the SDEM(GOx-HRP) can detect the 200 µM while free enzymes need 50 min reaction time. Those results provide support to the kinetics results in Figure 2c. Although at 50 min reaction time, both free enzymes and SDEM(GOx-HRP) system can detect down to 200 µM GO, we calculate the inferred GO concentration accurately by the standard curves R^2^ in Figure 3i. The SDEM system has bigger R^2^ with smaller error bar than free enzymes.

The SDEMs system can be extended to other enzymes cascade reactions like SDEM(SOX-HRP) (Supplementary section S5). The detection performances are based on the enzyme activity of input enzyme concentration (Fig. S5-2). For SDEM(SOX-HRP) system, it has a LOD of 16 µM while free enzymes system has a LOD of 40 µM with 20 nM SOX and 20 nM HRP (Fig. S5-3).

### Tunable and dynamic SDEMs on demand with different cycles

DNA molecules can be constructed into different tunable structures based on hybridization energy. We used a DNA hairpin to connect GOx and HRP to construct a SDEM(GOx-HRP), which is called a close status. A trigger strand containing 5, 2*, 3*, and 2 can bind to the hairpin structure of 2*, 3, and 2 to make this close status to an open status (see Fig. 4a). The open status has a toehold end of 5 which can be displaced with displacement strand so that the open status will be recycled into close status. We can add more trigger strands and displacement strands to make the close and open status cycling. Moreover, the detection performance can be changed by the close-open status and different cycling on demand. The signal of open status is always bigger than close status in the same cycle (Fig. 4b). We also validated the cycling processes by fluorescence change in Figure 4c. In order to expand the programable assembly, we added one more hairpin structure (Fig. 4d). The cycling performances have been summarized in Figure 4e. Notably, cycle 2 has the biggest *Δ*Signal (open – close) in one hairpin or two hairpins (Fig. 4e). The strand displacement was verified by PAGE gel in Figure 4f. In some cases, we can add three hairpins in SDEM(GOx-HRP) on demand (Fig. 4g-h). The SDEM(GOx-HRP) yields with hairpin structures were also simulated with DNA concentrations, temperatures and binding energies (Fig. 4i). Under our enzyme reaction temperature of 25 °C, the yields with designed structures of −25 kcal mol^-1^ are all 100% which indicate a stable structure. The detail of cycling programable assembly is shown in the section of Supplement 6 (Fig. S6-1 to Fig. S6-3).

**Figure 4.**
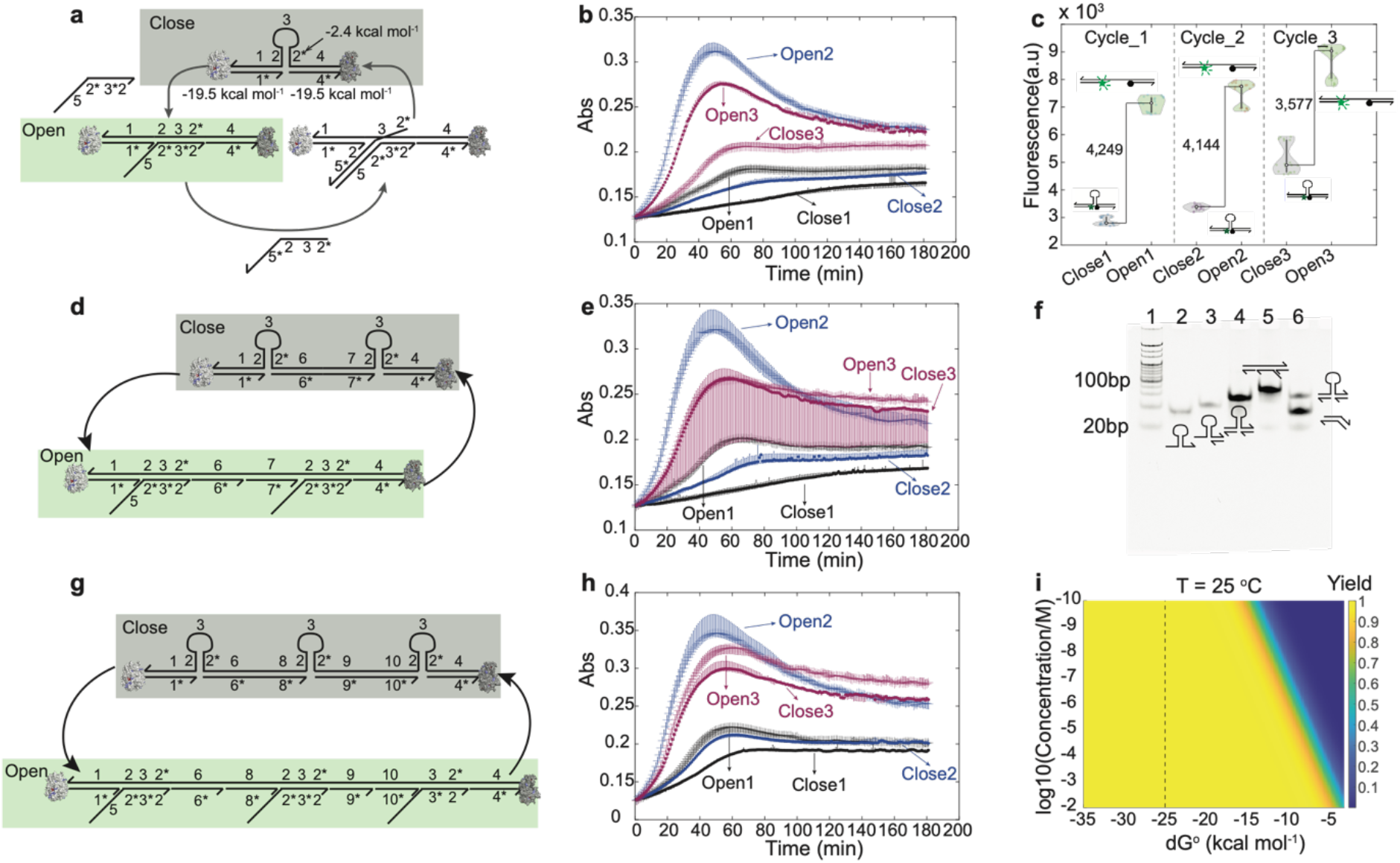
Programmable SDEMs on demand with multiple cycles. a. Toehold DNA strand triggered programmable SDEM change on demand. Number labels denote functional domains, which are continuously stretches of DNA that act as units in binding. Domain x* is the complement of domain x. b. The enzymes catalytic activity of SDEM with 3 cycles of close and open status. Final each enzyme concentration is 1 nM and concentration of GO is 4 mM at 37 °C. c. Demonstration of programmable SDEM in cycling. The carboxyfluorescein (FAM) is labeled in 3’ end of 1* and BHQ1 is labeled in 5’ end of 4*. The fluorescent reporter can be quenched and emitted with different cycles on demand. d. Programmable SDEM on demand with two hairpins strategy. Triggering open strand and triggering close strand can trigger the cycles from open to close. e. Enzymes catalytic activity of two hairpins SDEM with 3 cycles on demand. The dots show the median of triplicate and lines show the max and min values of the Abs. f. Verification of the programmable SDEM dynamic on demand of a. The structures of products of a are quantified with 8% PAGE gel at 80 V for 2 h. g. Programmable SDEM on demand with three hairpins strategy. h. The enzymes activity of programmable of SDEMs on demand of 3 cycles. i. The hybridization yield simulation of different hybridization thermodynamic energy.

### Catalyst assemble SDEMs on demand

Besides programmatically controllable in configuration, the SDEMs can also form a new SDEM by just adding a catalytic DNA strand. David Yu Zhang et al reported an entropy-driven reaction catalyzed by DNA and the reaction is simple, fast, modular, composable, and robust^41^. The scheme was shown in Figure 5a. This entropy reaction yield is as high as a directly annealed reaction and validated by PAGE gel without enzyme assembly (Fig. S7-1). The catalytic assembly yield was quantified by detecting 6-carboxy-X-rhodamine (ROX) which released from strand a-b using fluorescence spectroscopy (Fig. S7-2). We applied this entropy-driven reaction to catalyze one SDEM(GOx-HRP) to a new SDEM(SOX-HRP) (Fig. 5a). SDEM(GOx-HRP) is constructed by HRP and GOx via three DNA strands of A, B, and C. Catalytic strand (D) can displace strand B since C has the toehold of e*. The reaction *Δ*G can be calculated in equation 1 which is about −2 kcal mol^-1^. After the reaction, strand B with GOx will be released. By adding strand E with SOX, the second strand displacement reaction will help form SDEM(SOX-HRP). This reaction *Δ*G is also −2 kcal mol^-1^ by calculating with equations 5-8.

**Figure 5.**
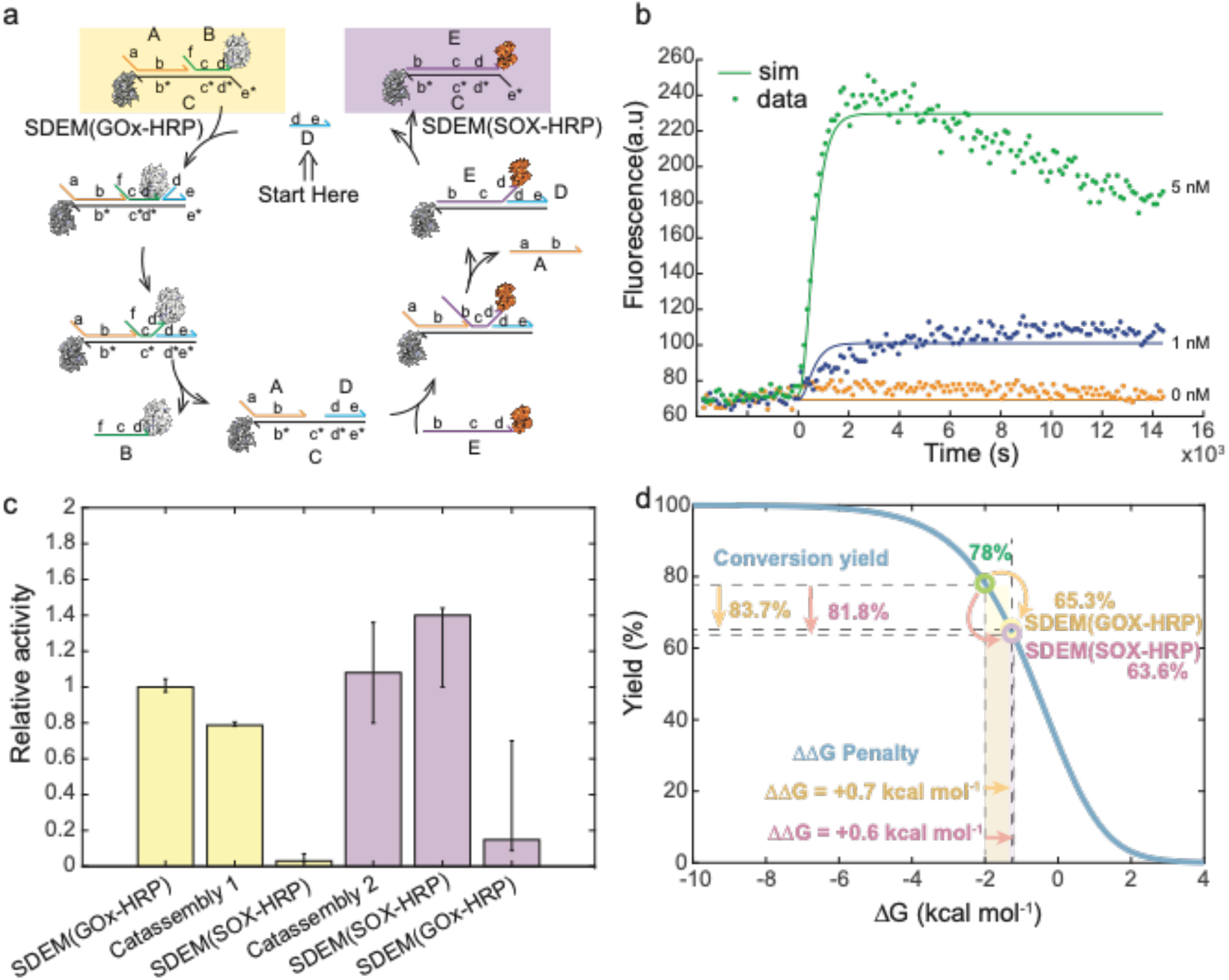
The catalytic assembly of SDEM(GOx-HRP) to SDEM(SOX-HRP). a. Schematic of catalytic assembly of SDEM(GOx-HRP) to SDEM(SOX-HRP). b. Demonstration of catassembly of SDEM. Different amounts of catalyst were introduced into the system at t = 0. The control trace (orange) shows the reaction with no substrate and no catalyst. The blue trace and green trace were investigated with 1 nM and 5 nM catalyst strand. c. Relative activity of before and after the catassembly. The box data (yellow) in triplicate are of detecting GO with three states. Catassembly 1 is the state of SDEM(GOx-HRP) after strand B displacement which is a mixture of B and ACD in figure 5a. The data (purple) are performed in triplicate of each box for sarcosine detection. Catassembly 2 is a mixture of ACD and E in figure 5a. The data (purple) of SDEM(GOx-HRP) system were detected without SOX. d. The influence of enzyme conjugation to catalytic assembly.

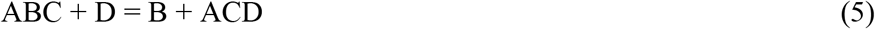

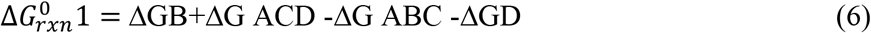

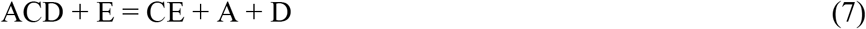

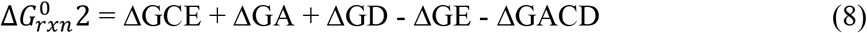

The rate constants k_0_, k_1_, k_2_, k_3_, and k_ROX_ were measured and simulated based on the data of Fig. 5c (equations 9-12). The catalytic reaction time over a wide range of catalyst concentrations is accurately reproduced by this reduced system of rate equations (Fig. 5b). According to Zhang’s model^41^, the addition of catalyst can accelerate the reaction by over six orders of magnitude (k_2_/k_0_ =2.3*10^6^).

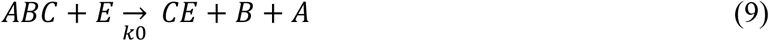

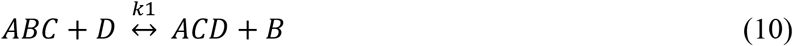

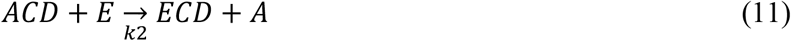

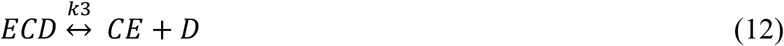

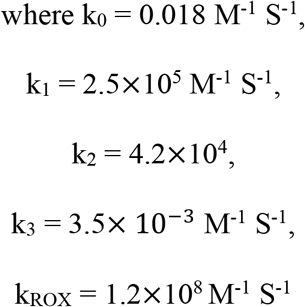

We also quantified SDEMs enzyme activities before and after catalyst assembling in Fig. 5c (Fig. S7-3 to Fig. S7-6). After 2h catalytic assembly displacement by strand D, the GOx was released from SDEM(GOx-HRP). The relative activity decreased by 26%. When the catalytic assembling was finished, the activity of SDEM(SOX-HRP) was decreased by 97%. This decrease may result from two parts, one is additional SOX diluting the GOx-HRP cascade reaction, the other is that the domain HRP activity was taken by SDEM(SOX-HRP). Since the SOX-HRP was from free state to assembly state, the activity of SOX-HRP cascade went higher. We calculate the *ΔΔ*G using the following equations:

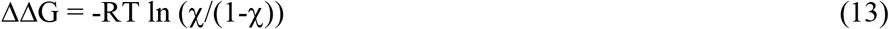

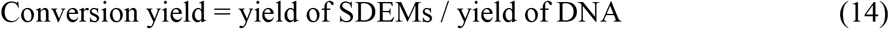

In order to quantify the yield of the catalytic assembly, we calculate the theoretical yield of 78% based on *ΔΔ*G in equation 13 (*ΔΔ*G = −2 kcal mol^-1^). The real yields of SDEM(GOx-HRP) and SDEM(SOX-HRP) are 65.3% and 63.6% respectively (Table S7-1 and Table S7-2). The conversion yields of SDEMs were calculated using equation 14 and are shown in Fig. 5d. Those constants were used for quantifying the enzyme’s influence in the catalytic assembly. The thermodynamics energies change of adding enzymes was calculated by equation 13. Consequently, the *ΔΔ*G penalties of GOx and SOX are +0.7 kcal mol^-1^ and +0.6 kcal mol^-1^ respectively which are minor changes since *ΔΔ*G of one base change is about ± 2 kcal mol^-1^.

## Discussion

In this study, the DNA-enzyme super molecules presented here notably achieve programmable cycling and catalytic assembly on demand. This method may mimic the natural metabolism, controllable biosynthesis, and low concentration of metabolons detection. Although enzyme engineering can control enzymes cascades, we are not aware of any other technologies that can programmatically control enzymes cascade cycling and catalytic assembly. Some DNA nanostructures can control cascade enzymes assembling^9,10,31^ but the DNA nanostructures are complex and comprised of multiple DNA strands with negative charge which will influence the enzyme’s activities^40,42^. Generally speaking, other enzyme engineering technologies are fixed assembly which are not flexible enough to do programable cycling and catalytic assembly on demand.

In this work, we first demonstrated two SDEMs with two enzymes cascades that can do programable cycling and catalytic assembly. We applied catalyst strand to programable SDEMs technology. This catalyst reaction can accelerate the reaction by over six orders of magnitude. We also noticed the enzyme which linked to the DNA strand contributes little to the thermodynamic changes of DNA strand displacement and the energy change is less than the energy from one binding nucleotide. This minor change can be ignored during SDEM designs and it also shows the proof of the thermodynamic calculation of DNA-enzyme conjugation system. In our preliminary data, the SDEM catalytic activity is related to the enzymes’ intrinsic activities, like SDEM(GOx-HRP) and SDEM(SOX-HRP) having a similar yield of about 64% but activity differences are huge. However, this new catalytic assembly provides a new solution to the detection of substrates on demand. We can use SDEM(GOx-HRP) to detect glucose, and we just need to add catalytic strand to form an SDEM(SOX-HRP) if we want to switch to detect sarcosine. This will make the detection on demand more flexible.

To date, most enzymes cascade assembles depend on either DNA nanostructures or peptide scaffolds (Table 1). The DNA nanostructures include DNA origami, DNA tetrahedral, and DNA scaffolds need a lot of DNA strands which will be cost consumable. We estimate the $0.2 / nt so that the cost for DNA origami will be about $2900 while our method only needs about $20 which is 100-fold lower than the cost of DNA origami. For the enzymes assembly which depends on peptide, is hard to be made programmable for its high binding efficiency. At the same time, the assembly with toehold displacement provides a new solution of programable with low cost. Previous works all used toehold to archive the enzymes assembly and disassembly in one system with two enzymes. However, the in vivo metabolic pathway is complicated. Here, we also expand to catalytic assembly from one SDEM(GOx-HRP) to another SDEM(SOX-HRP) by just adding a DNA catalyst strand.

**Table 1.**
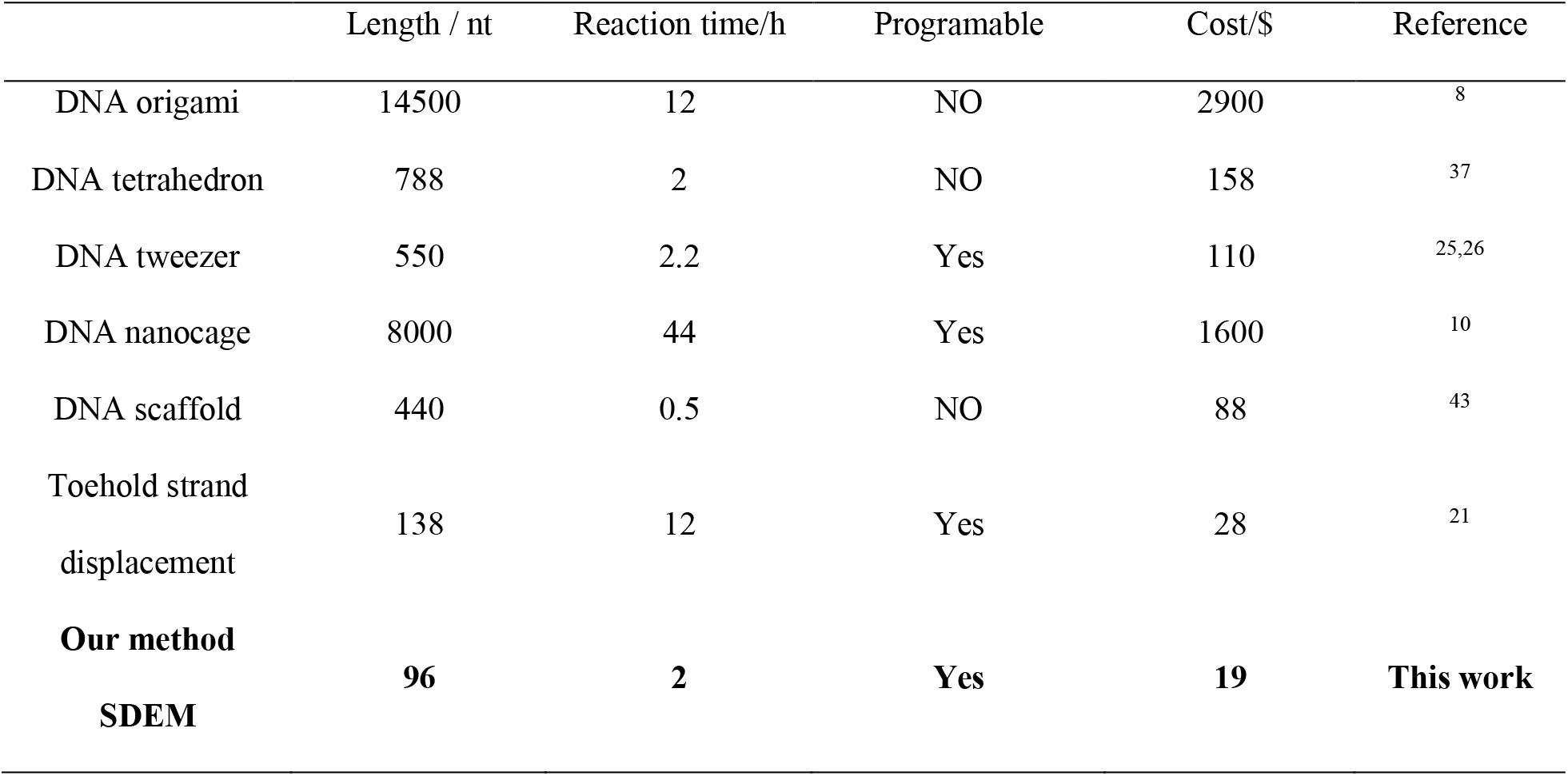
Comparison of the performance of different enzymes assembly methods.

In this present design, the SDEM technology is primarily introduced to the enzymes’ cascades which are sensitive to distances. Thus, the SDEM is well-suited for the detection of actionable low concentration of metabolons with low cascade enzymes. For the system with high enzymes concentration, the distances between enzymes are small and the distance changes of programable SDEMs are too minor to influence the enzymes activities. However, it is hard to reach a very high enzymes concentration in vivo and high-concentration dependent distance control is not controllable and programable. Moreover, the conjugation position will influence the activity if in the enzymes’ activity centers. The potential solution may need to take advantage of non-modification enzyme technologies like the enzyme binding aptamer.

In summary, we constructed a SDEM to provide a new solution of programable controlling enzymes cascades activities on demand. Moreover, through adding one catalytic DNA strand, we achieved different SDEMs conversions on demand to sense different metabolisms like GO and sarcosine. Given DNA strand displacement and enzyme DNA high-yield conjugation, we do think this method has multiple potentials to be applied in other enzymes assembly on demand since most enzymes have lysine residues which are linkage points to DNA. On the other hand, programable reactions depend on DNA strand displacement which are on account of DNA hybridization thermodynamics and can achieve controlled reactions of fast rate or slow rate. Besides two enzymes’ cascades, we think this method can be expanded to three, four or more enzymes’ cascades to mimic more complex in vivo metabolism reactions in vivo.

## Methods

### Materials

GOx (G7141) from *Aspergillus niger* (type X-S) HRP (P6782), D-glucose N-succinimidyl 3-(2-pyridyldithio) propionate (SPDP) and dimethyl sulfoxide (DMSO) were purchased from Sigma-Aldrich, USA. 3,3’,5,5’-tetramethylbenzidine (TMB, 34021) were purchased from ThermoFisher Scientific, Inc., USA. SOX (60105) was purchased from Realbio (Shanghai, China). All DNA probes were synthesized and modified by Sangon Biotech. The other chemical reagents were purchased from Sinopharm Chemical Reagent Co. Ltd. (Shanghai, China). Milli-Q water from a Millipore system was used throughout all the experiments for all solutions preparation.

### Enzyme-DNA conjugation

GOx was modified with linker1 (L1) and HRP was modified with linker2 (L2). An SPDP molecule was used to link an oligonucleotide with an enzyme. Firstly, the GOx was incubated with the SPDP (5-fold excess, 100-fold excess for HRP-DNA conjugation) for two hours in Dulbecco’s Phosphate Buffered Saline (DPBS) (pH 8), which resulted in an SPDP to link with the lysine residues of the enzymes with amine-reactive N-hydroxysuccinimide (NHS). The SPDP is diluted with DMSO. The excessive SPDP was removed by washing with DPBS, using 30 kD cutoff filters. Secondly, the SPDP-enzyme was incubated with thiol-modified DNA (10-fold excess) and the thiol-modified DNA was activated with a 20-fold tris(2-carboxyethyl) phosphine hydrochloride (TCEP). Lastly, the excessive DNA was removed by washing using the 50 mM HEPES (1 M NaCl, pH=8). All the enzymes were quantified by UV-visible light spectrophotometer (Lambda850, USA).

### SDEM(GOx-HRP) assembly

1 μM GOx-L1 and 1 μM HRP-L2 were incubated with 1 μM L3 in TM buffer (20 mM Tris, 50 mM MgCl_2_, pH 8.0) at 37 °C for 2 hours. And then the GOx-HRP conjugation was stored at 4 °C.

### AFM imaging

1‰ (3-aminopropyl)triethoxysilane (APTES) was incubated on freshly cleaved mica surface for 30 s, and washed off by Q water. 20 nM SDEM (in TM buffer) were spotted onto mica surface which is modified with APTES for 1.5 min to allow the enzymes to absorb the substrate and then washed off by Q water. The assembled enzyme pairs were characterized by AFM (Multimode Nanoscope VIII, Bruker) in ScanAsyst mode. The yield of assemblies with two enzymes of the SDEM(GOx-HRP) was ∼47.6% as shown in Figure S2-4. The assembled enzyme pairs appear as individuals with a distance between 10-20 nm, which was in agreement with the designed structure. The free enzymes and SSDMEs(GOx, HRP) were immobilized with the same method for AFM imaging.

### TEM characterization

The enzyme-DNA conjugation was prepared via SPDP method in Section S1. The linker DNA L3 was added into the enzyme-DNA conjugation (GOx-SPDP-L1, HRP-SPDP-L2) with L3 to the enzyme-DNA conjugation molar ratio of 1:1 (GOx-SPDP-L1) and molar ratio of 1:1 (HRP-SPDP-L2), respectively. The SDEM(GOx-HRP) were directly characterized with TEM (120 kV; Talos L120C G2) as shown in Figure S2-8. Statistical results of the SDEM (GOx-HRP) showed that the yield of assemblies with two enzymes of the SDEM(GOx-HRP) was ∼52.2% as shown in Table S2-1.

### SDEM cascade assays in solution

The SDEM(GOx-HRP) cascade assays were conducted with the SpectraMax iD5 96 well plate reader (Molecular Devices, USA). 0.064 g/L TMB substrate was added for monitoring the absorbance change at 370 nm in 4 mM glucose (pH 7.4). The final concentration of SDEM(GOx-HRP) was 0.8 nM as shown in Figure 2c. The free enzymes and SSDMEs(GOx, HRP) were in the same condition for enzyme cascade assays.

## Code availability

The MATLAB and R codes used for data analysis and simulation are available upon request under NDA for academic lab.

## Data availability

All data supporting the findings of this study are available from the corresponding author upon reasonable request.

## Acknowledgements

This work was financially supported by the Ministry of Science and Technology of China (2021YFF1200302), NSCFC (22174094).

## Author contributions

H.P.Z., X.L.Z., C.H.F., and P.S. conceived the project. H.P.Z., L.Y.T., G.M.Y., and P.S. designed and performed the experiments. H.P.Z., X.X.H., L.M.Q., and P.S. analyzed and simulated the enzyme cascade activity data. H.P.Z. and P.S. wrote the manuscript with input from all authors.

## Additional information

Supplementary information: The online version contains supplementary material available.

