## Supplemental information for "Programming super DNA-enzyme molecules for on-demand enzyme activity modulation"

**S1. DNA-Enzyme Conjugation.**

**S2.** **Assembly and Characterization of the SDEM.**

**S3. The Optimization of SDEM Cascade Activity.**

**S4. Nonspecific DNA Interactions.**

**S5. Verifying the Enhancement SDEM Cascade Reactions for Sarcosine Detection.**

**S6: The Programable SDEM.**

**S7: The SDEM Cascade Reactions and Networks Catalyzed by DNA.**

**S8. DNA sequencies.**

**Section S1: DNA-Enzyme Conjugation**

**Three-Dimensional structures and the lysine residues of enzymes.** Three-dimensional structures of enzymes from PDB (www.rcsb.org) were shown in Figure S1-1 via PYMOL. The distribution of enzyme particles’ sizes in the XY-axis was labeled and the sizes were obtained by calculating the boundary distance of atoms in the xy axis in PYMOL, GOx (x = 5.0 nm, y = 7.2 nm), HRP (x = 4.0 nm, y = 6.2 nm) and SOX (x = 5.9 nm, y = 8.8 nm). The molecular weights of the enzymes were 65.7 kD (GOx), 34.8 kD (HRP) and 88.0 kD (SOX), respectively, which were obtained from the website (www.sigmaaldrich.cn). The lysine residues of the enzymes on the enzyme surface determined the SPDP-to-enzyme ratio. The total number of lysine residues and the number of lysine residues on the enzyme surface were shown in Table S1-1. The surface residues of the enzyme to the total number of lysine residues of the enzyme were the ratio of the surface lysine residues.

**SPDP-Enzyme conjugation**. The enzyme and DNA were linked by SPDP^1^. The amine-reactive N-hydroxysuccinimide (NHS) esters of SPDP reacted with the primary amines of lysine residues on the enzyme surface (pH 8.0) as shown in Figure S1-2^2-4^. In this reaction, the yield of SPDP-enzyme was sensitive to the pH value^5^. For a neutral or acidic pH, the primary amines were protonated and reacted slowly with the NHS group of SPDP, and therefore, this reaction was in alkaline pH. The primary amines of amino acids residues exist in lysine, asparagine, glutamine and arginine in twenty amino acids as shown in Figure S1-3(a-d). In the presence of base, the primary amines of asparagine, glutamine and arginine have resonance structures of an amide group and have no ability to react with the NHS group of SPDP as shown in Figure S1-3(e). Thus, only the primary amines of lysine residues reacted with the NHS group of SPDP by nucleophilic displacement with a primary amine, generating a new amide bond as shown in Figure S1-3(f) and Figure S1-2(1). The surface lysine ratio was shown in Table S1-1 and the high surface lysine ratio contributes to the reaction between SPDP and the primary amines. A thiol-modified oligonucleotide was conjugated to the SPDP-modified enzyme through the disulfide bond exchange between the thiol-modified oligonucleotide and SPDP as shown in Figure S1-2(2).

**Quantification of DNA-enzyme conjugation efficiency via absorbance spectra.** The thiol-modified oligonucleotides were added to the solution of SPDP-modified enzymes and pyridine-2-thione (ε=8080 M^-1^ cm^-1^ at 343 nm) was then released. The concentration of GOx, HRP and SOX were quantified by measuring the absorbance at 452 nm, 403 nm and 452 nm, respectively. The DNA labeled on the surface of enzyme were quantified by measuring the absorbance of pyridine-2-thione at 343 nm. The labeling ratio of DNA molecules was estimated using equation 1 as shown below:

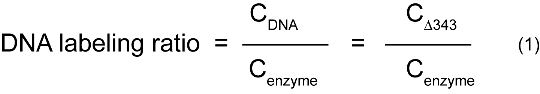

An average DNA-to-enzyme ratio was obtained and the labeling ration of DNA molecules to the GOx, HRP and SOX were 2.2, 1.4 and 1.4 respectively, as shown in Figure S1-4.

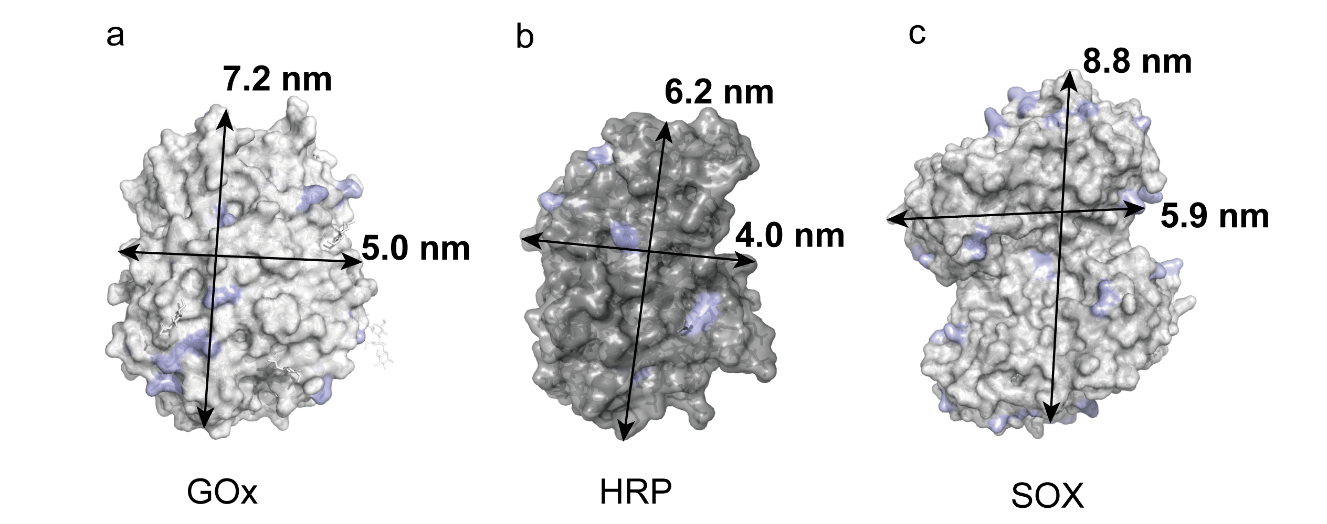

**Figure S1-1.** Three-dimensional structures of enzymes from PDB. (a) GOx (65.7 kD, PDB code: 1CF3) (b) HRP (34.8 kD, PDB code: 1H5A) (c) SOX (88.0 kD, PDB code: 3QSE). Lysine residues were shown in light blue.

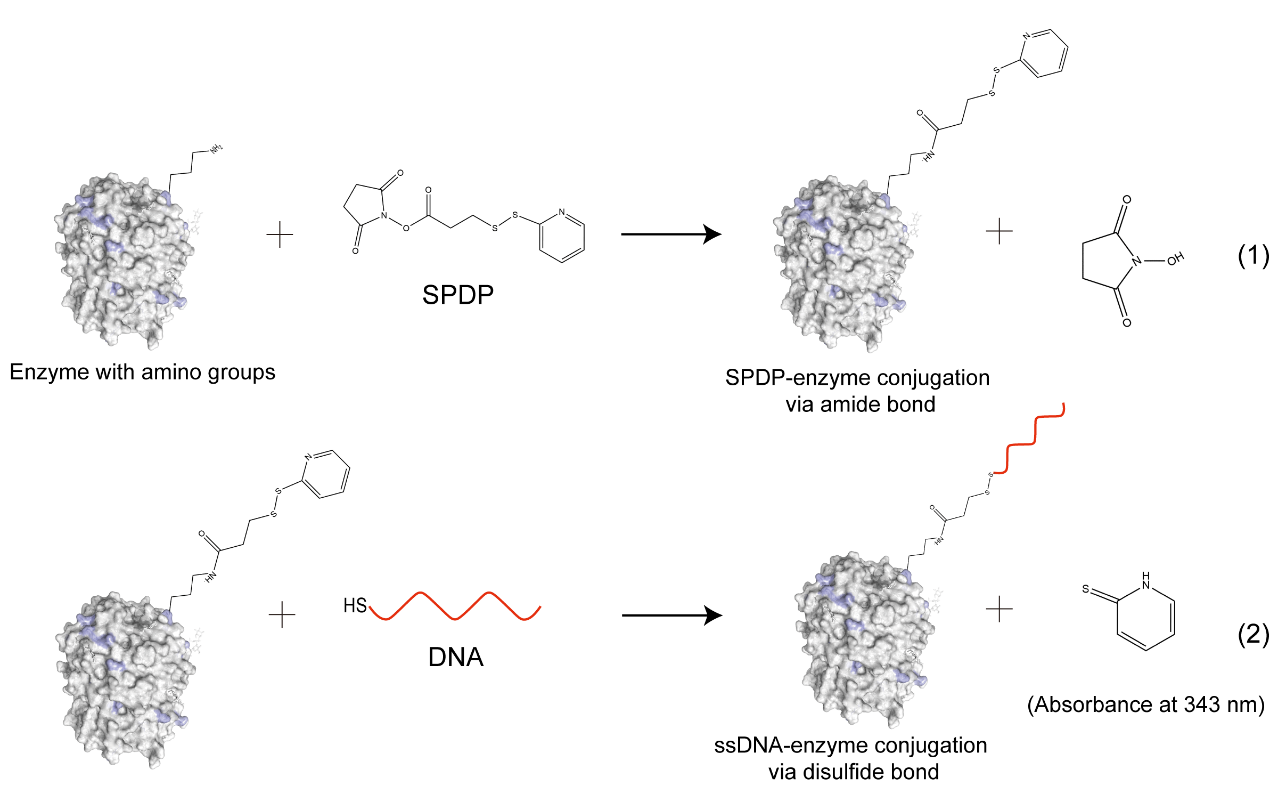

**Figure S1-2.** DNA-Enzyme conjugation using a SPDP crosslinker.

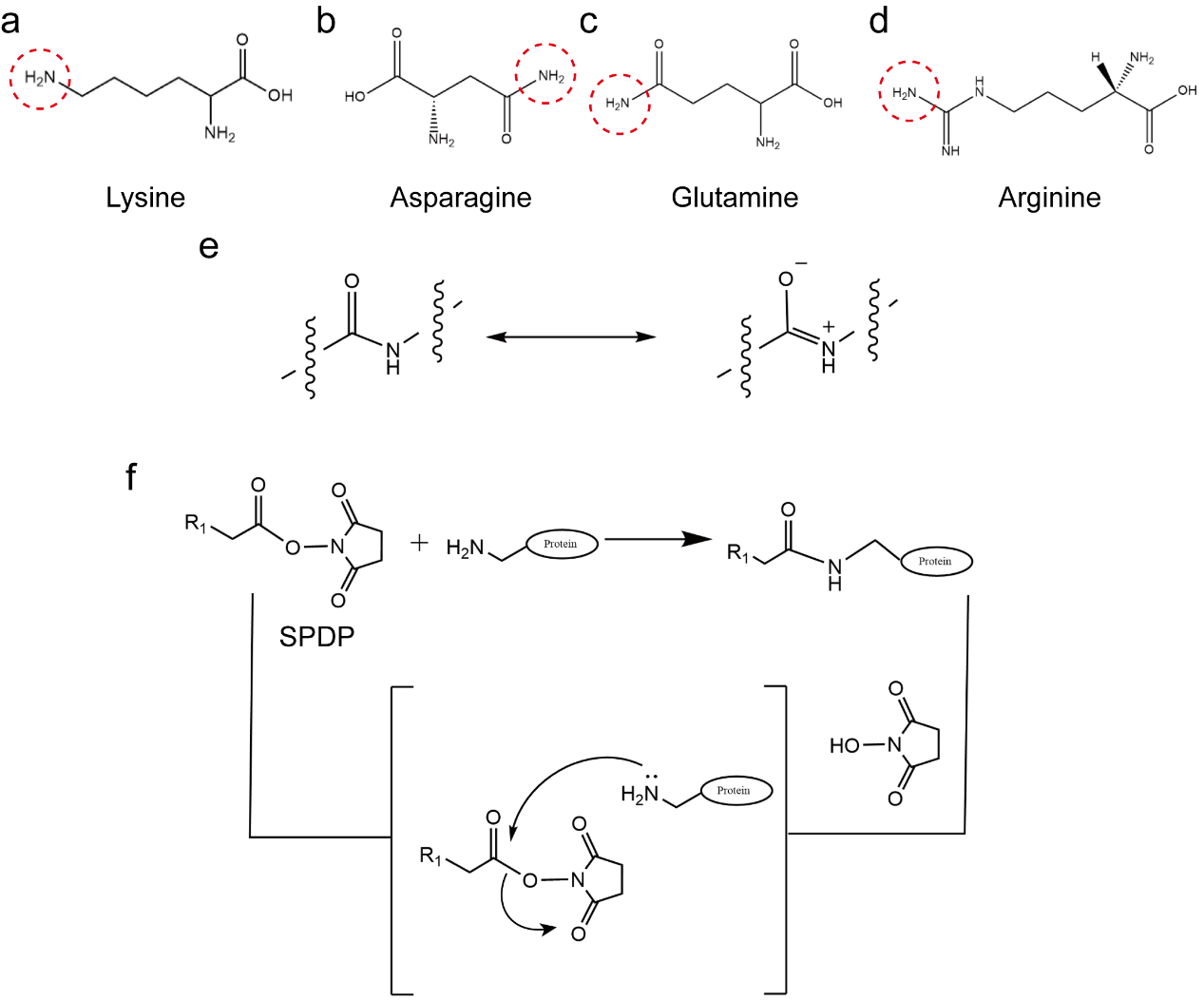

**Figure S1-3.** The principle of the reaction between lysine on enzymes and SPDP. (a) Lysine (b) Asparagine (c) Glutamine (d) Arginine. The primary amines of amino acids residues were in red-dashed circles. (e) Resonance structures of an amide group. (f) The formation of a new amide bond between lysine on enzymes and SPDP. R_1_ is the complementary part of the SPDP molecule.

**Table S1-1.** The total lysines and the surface lysines of GOx, HRP and SOX.

| Enzymes | Total lysine | Surface lysine | Surface lysine ratio |
| --- | --- | --- | --- |
| GOx | 15 | 15 | 100.0% |
| HRP | 6 | 6 | 100.0% |
| SOX | 42 | 38 | 90.5% |

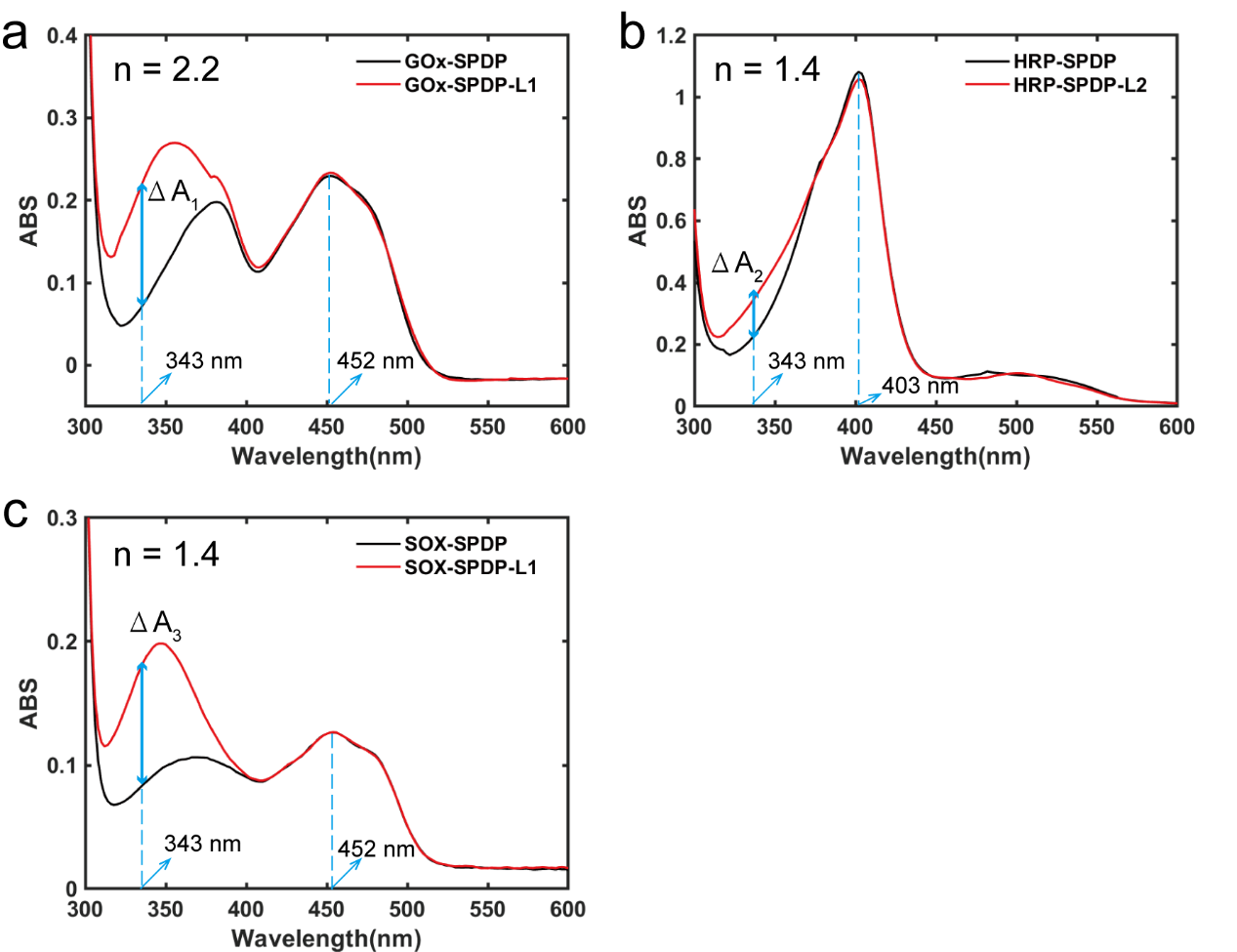

**Figure S1-4.** Quantification of DNA-enzyme conjugation efficiency via absorbance spectra.

(a) GOx-L1 conjugation: ∆A_1_ at 343 nm before and after L1 conjugation is ~ 0.2 (extinction coefficient: 8080 M^-1^ cm^-1^), corresponding to 19.2 µM L1 coupled with 8.9 µM GOx (ε=28200 M^-1^ cm^-1^ at 452 nm for GOx). (b) HRP-L2 conjugation: ∆A_2_ at 343 before and after L2 conjugation is ~ 0.1 (extinction coefficient: 8080 M^-1^ cm^-1^), corresponding to 14.9 µM L2 coupled with 10.5 µM HRP (ε=100000 M^-1^ cm^-1^ at 403 nm for HRP). (c) SOX-L1 conjugation: ∆A_3_ at 343 before and after L1 conjugation is ~ 0.1 (extinction coefficient: 8080 M^-1^ cm^-1^), corresponding to 12 µM L1 coupled with 8.6 µM SOX (ε=12700 M^-1^ cm^-1^ at 452 nm for SOX).

**Section S2: Assembly and Characterization of the SDEM.**

**The preparation of the linked DNA strands of SDEM**. The paired DNA strands linked two enzymes of SDEM was three DNA paired strands (Fig. S2-1). All hybridizations were combined in equimolar quantities in TM buffer at 37 ℃ for 2 h and then the hybridization buffer was cooled to 4 ℃ in 30 s. The standard free energies of these DNA complexes were calculated at 37 ℃ (0.2 M Na^+^). The standard free energies of single strands were 0 kcal mol^-1^. The average free-energy change (ΔΔG) of L2+L3 was -22.5 kcal mol^-1^ in the **Figure S2-1** of reaction (1). The L1 was added to the solution of L2+L3 and the ΔΔG of the reaction (2) was -13.1 kcal mol^-1^. Therefore, L1 was prone to hybridize to the L2+L3. The assembled nanostructure can be used to link two enzymes for enzyme cascade reaction.

The construction of the assembled nanostructure was shown in **Figure S2-2** via NUPACK (www. nupack. org). The assembled nanostructure was synthesized with yields as high as 95.0% as shown in **Figure S2-3**.

**Preparation of assembled gold nanoparticles with double strand.** To proof the distance and linked yields of the DNA paired strands, we did TEM using gold nanoparticles linked DNA strands. The DNA modified gold nanoparticles (AuNPs) were prepared as reported methods^6^. The 5 nm AuNPs were modified with thiolated DNA B and thiolated DNA C, respectively. The two kinds of AuNPs were modified with different DNA were incubated with equal mole DNA solution (A2) at room temperature over 2 h. The assembled AuNPs were directly characterized with TEM (120 kV; Talos L120C G2) as shown in **Figure S2-4**.

**The SDEM preparation**. A double DNA strand nanostructure was used to link two enzymes to construct the SDEM, as shown in **Figure S2-5**. A double strand nanostructure and two kinds of enzymes were incubated in equimolar quantities in TM buffer at 37 ℃ for 2 h and then the hybridization buffer was cooled to 4 ℃ in 30 s. The inter-enzyme distance of the SDEM was as shown in **Figure S2-6(i)**.

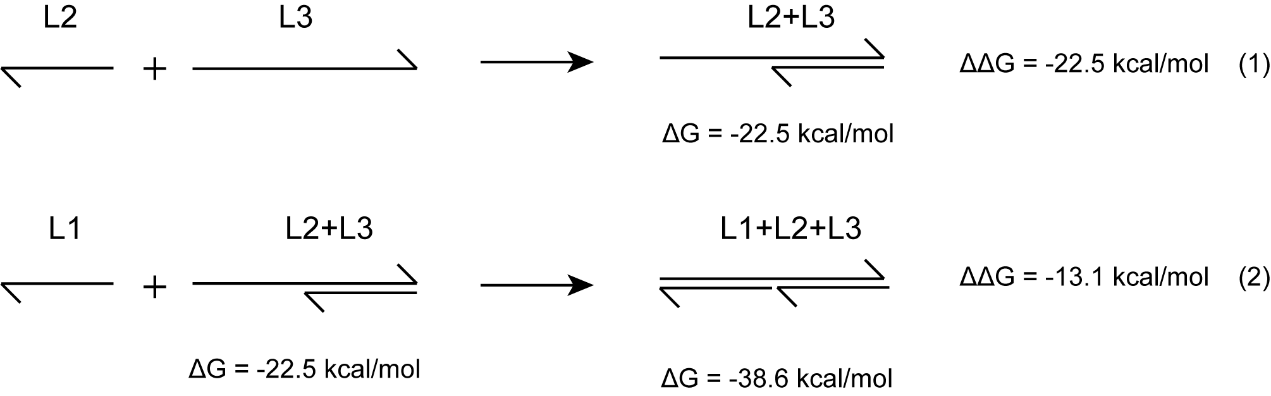

**Figure S2-1.** The ΔΔG calculation of the assembled nanostructure at 37oC.

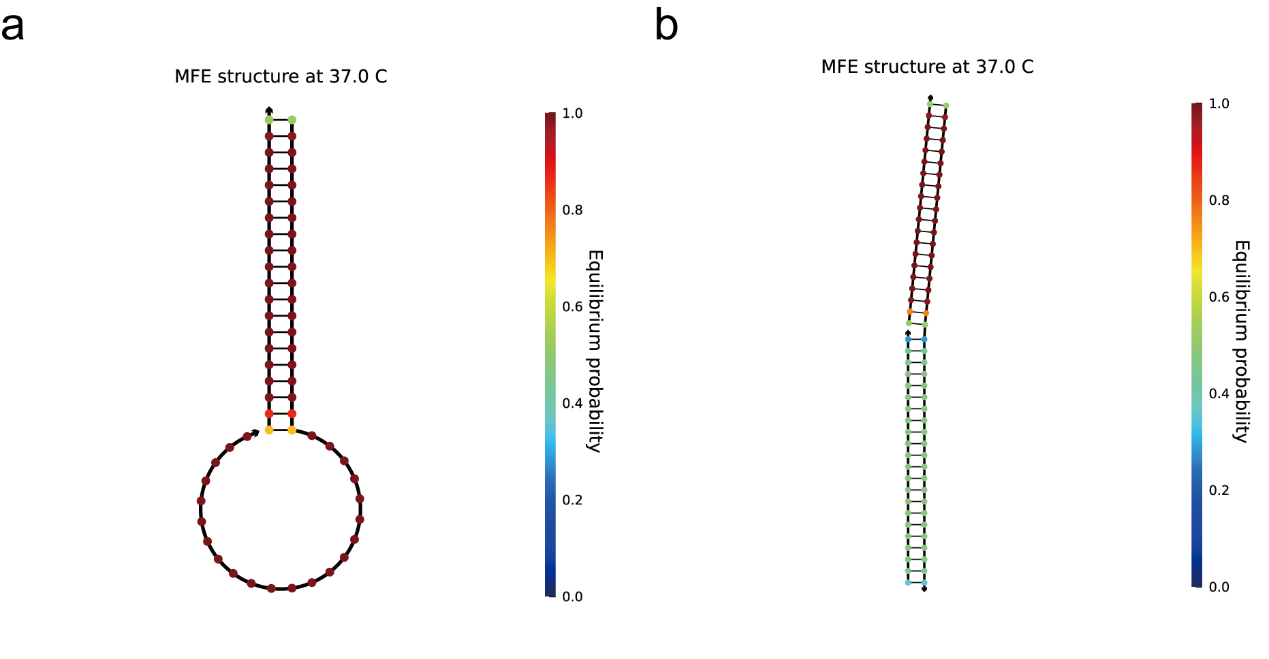

**Figure S2-2.** The secondary structures were predicted by the NUPACK software. (a) L2+L3 (b) L1+L2+L3

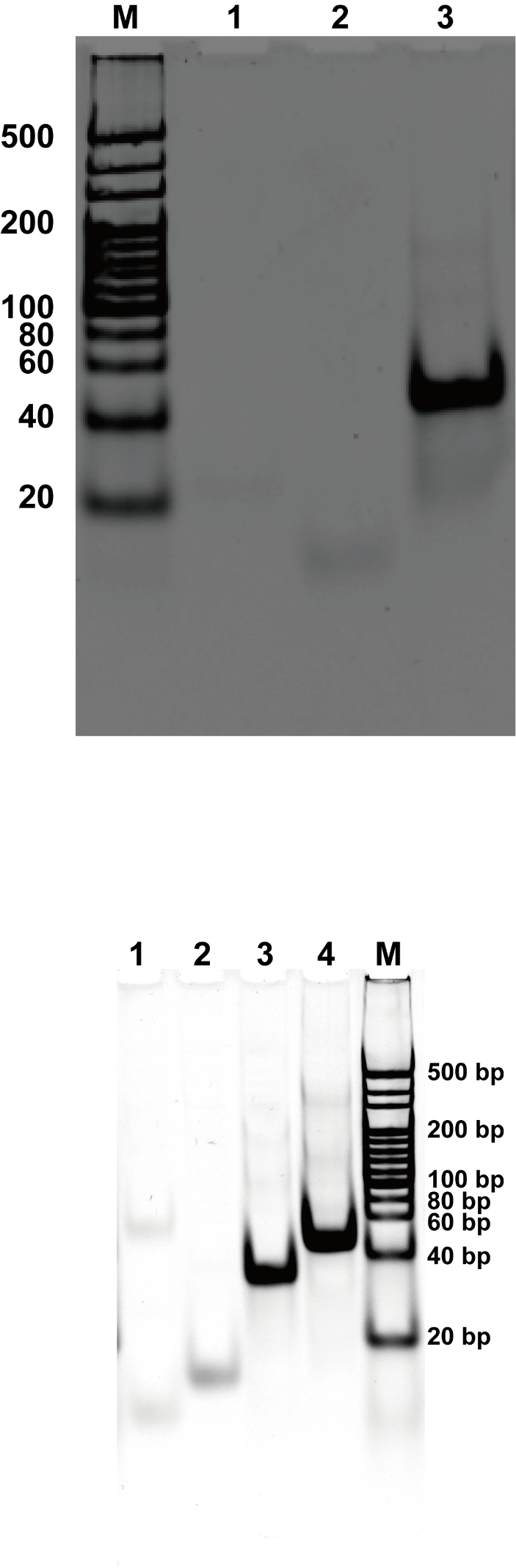

**Figure S2-3.** The construction of the assembled nanostructure. Native polyacrylamide gel electrophoresis of the assembled nanostructure. Lane 1: L2; lane 2: L1; lane 3: L2+L3; lane 4: L2+L3+L1; Lane 5: 20 bp DNA marker. Conditions: 12% PAGE with a constant voltage of 180 V for 1 h.

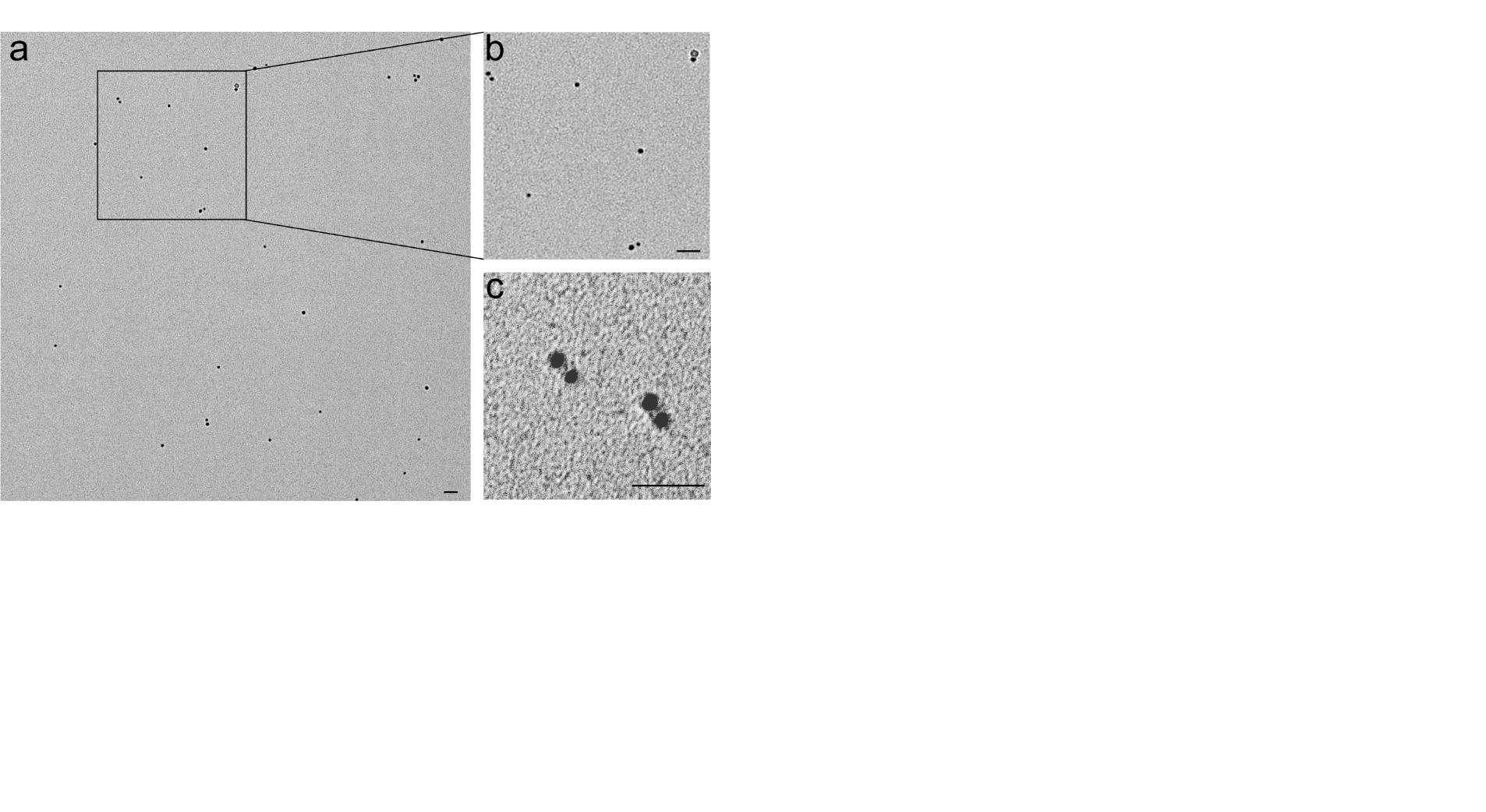

**Figure S2-4.** dsDNA-Directed assembly of AuNPs. (a) The TEM image of assembled AuNPs (5 nm). (b) (c) High resolution images of the assembled AuNPs. The average distance for the two linked AuNPs is ~10 nm. Scale bar: 50 nm.

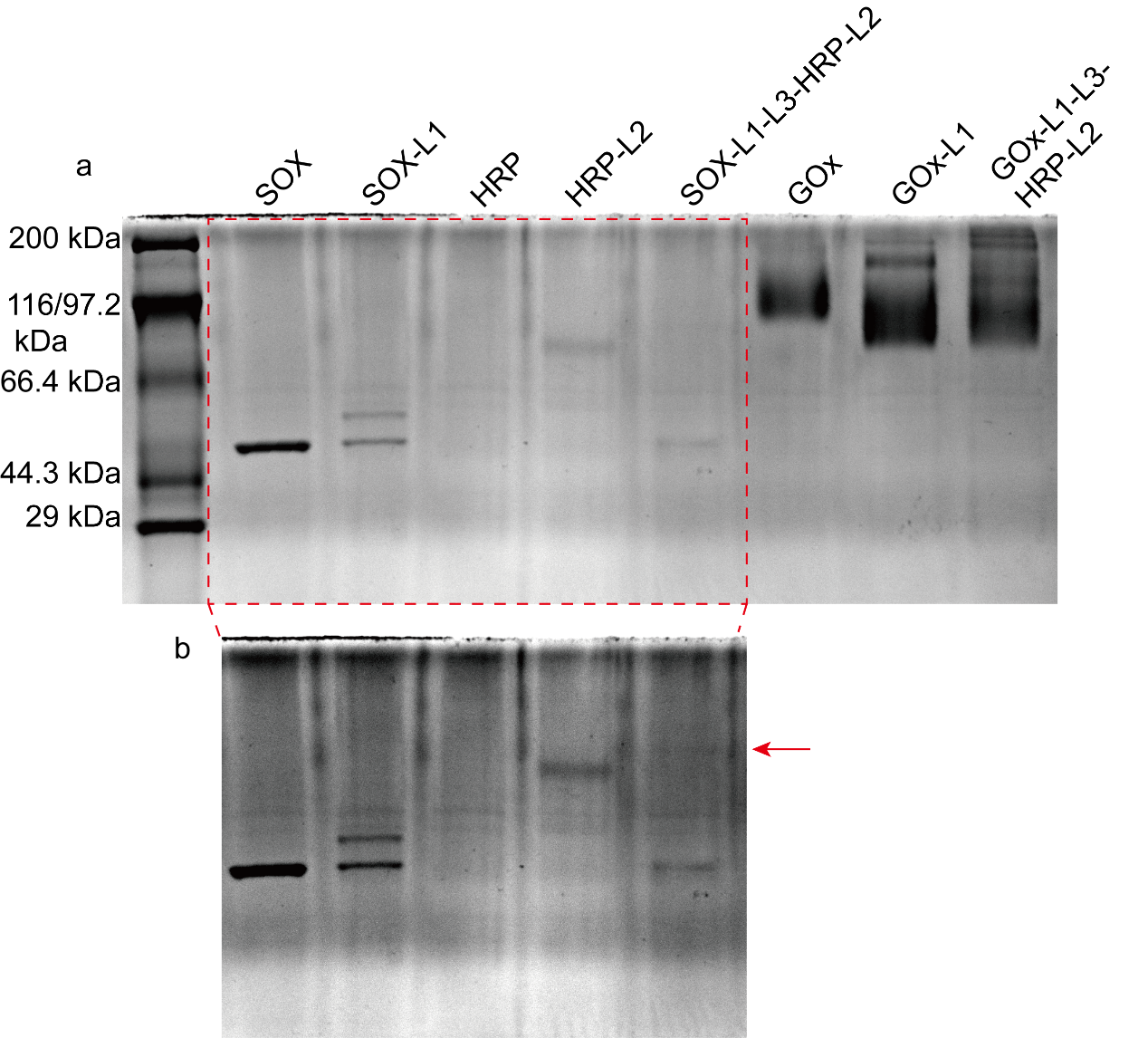

**Figure S2-5.** SDS-PAGE electrophoresis of protein-DNA conjugates and SDEM. (a) SDEM(SOX-HRP) and SDEM(GOx-HRP) assembly. (b) SDEM(SOX-HRP) assembly with higher gray value. Conditions: NuPAGE 5%-12% Bis Tris Gel with a constant voltage of 150 V for 60 mins.

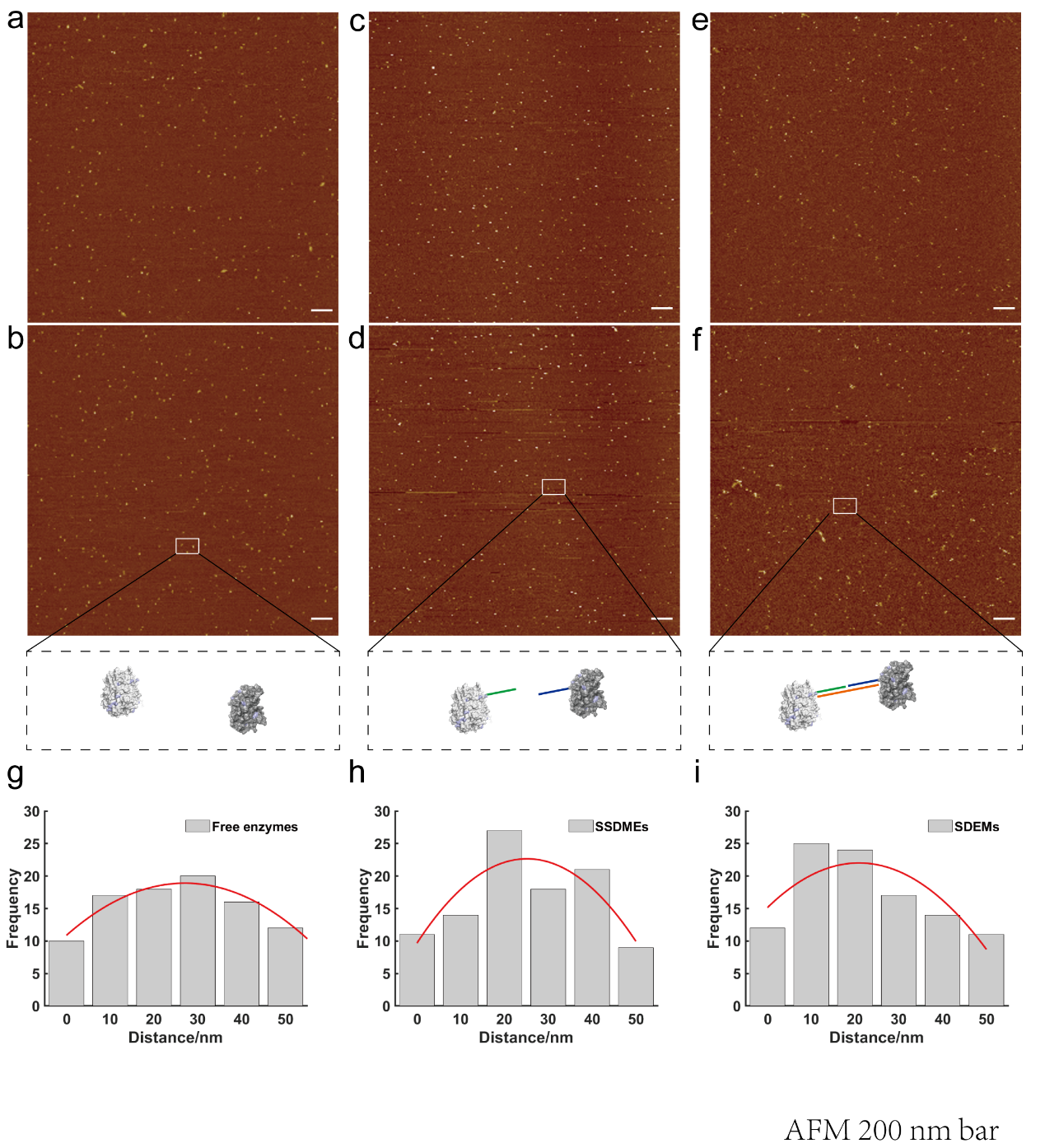

**Figure S2-6.** Evaluation the inter-enzyme distance of free enzymes, SSDMEs(GOx, HRP) and SDEM(GOx-HRP) on the mica. (a)(b) Free enzymes. (c)(d) SSDMEs(GOx, HRP). (e)(f) SDEM(GOx-HRP). (ghi) Statistical results of the inter-enzyme distance of free enzymes, SSDMEs(GOx, HRP) and the assembled enzyme pairs respectively. (g) Free enzymes. (h) SSDMEs(GOx, HRP). (i) SDEM(GOx-HRP). The inter-enzyme distance of the assembled enzyme pairs is 10-20 nm. Each AFM image is generated by scanning an area of 3 µm× 3 µm. Scale bar: 200 nm.

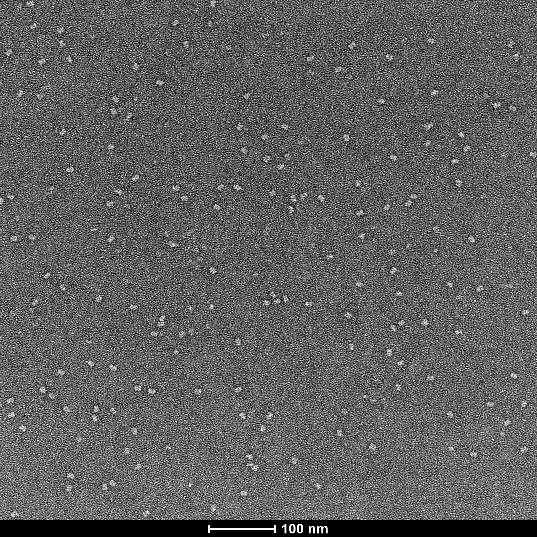

**Figure S2-7.** TEM image of the free enzymes. Different from SDEM(GOx-HRP), the particles of free enzymes were dispersed on the carbon film.

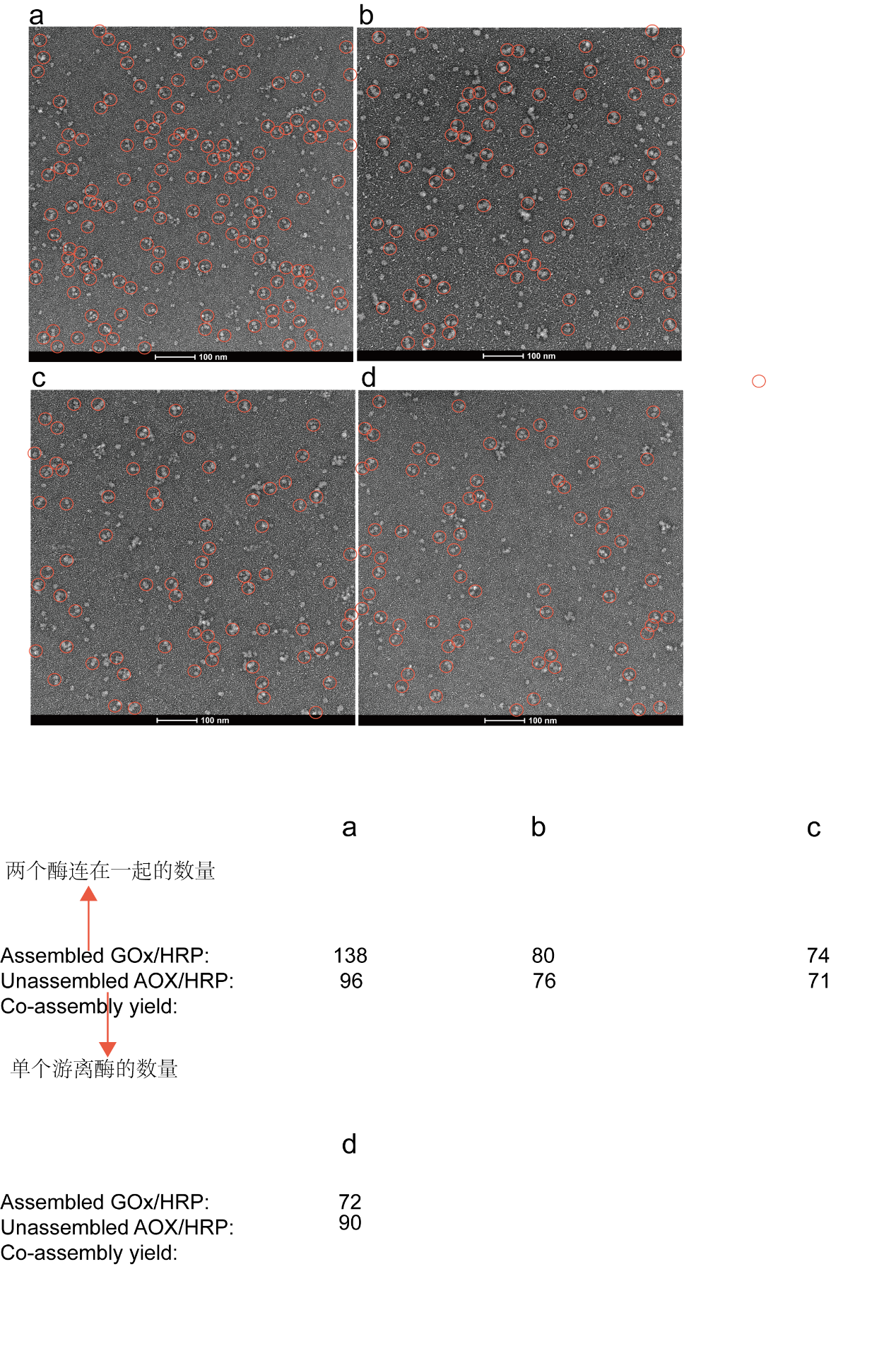

**Figure S2-8.** TEM images of the SDEM(GOx-HRP). dsDNA-directed assembly of GOx and HRP (a-d). The assembled enzymes were labeled with red circle. The assembled SDEM(GOx-HRP) had a yield of 52.2% and the statistics were as shown in Table S2-1.

**Table S2-1:** The yield of the assembled SDEM (GOx-HRP).

|  | **Assembled enzymes** | **Unassembled enzymes** |
| --- | --- | --- |
| **a** | 138 | 96 |
| **b** | 80 | 76 |
| **c** | 74 | 71 |
| **d** | 72 | 90 |
| **Total** | 364 | 333 |
| **Assembly yield** | 52.2% | |

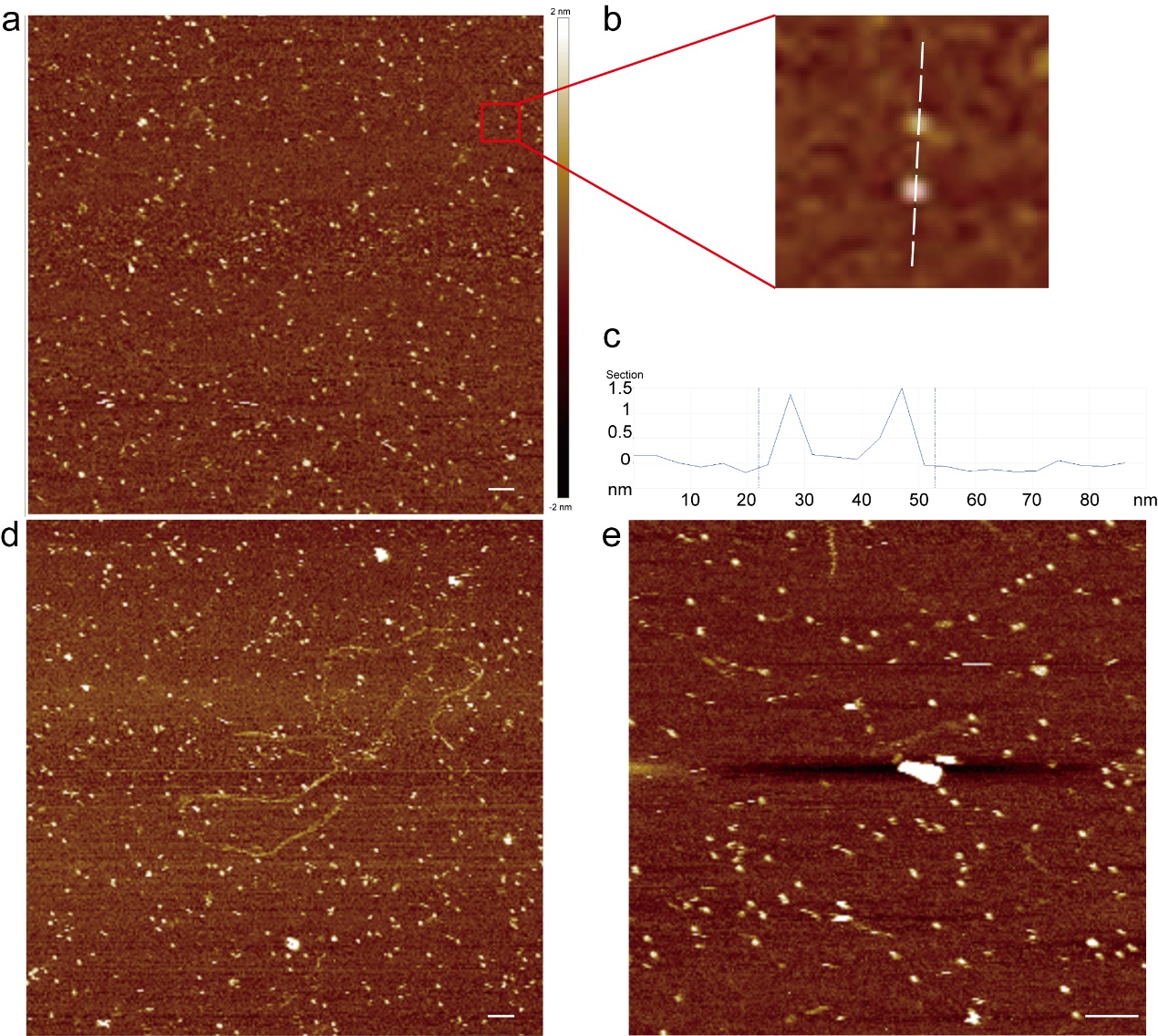

**Figure S2-9.** Characterization of the assembled SDEM(SOX-HRP) on the mica substrate (in 1× TM buffer). (a) AFM image of the assembled enzyme pairs linked with DNA double strand (42 bp). The colour bar indicated the height of the scanned surface. (b) High resolution images of the assembled enzymes from (a). (c) The height of the assembled enzyme pairs as shown in dash line from (b). (d)(e) AFM image of the assembled enzyme pairs at the same condition in (a). Scale bar: 50 nm.

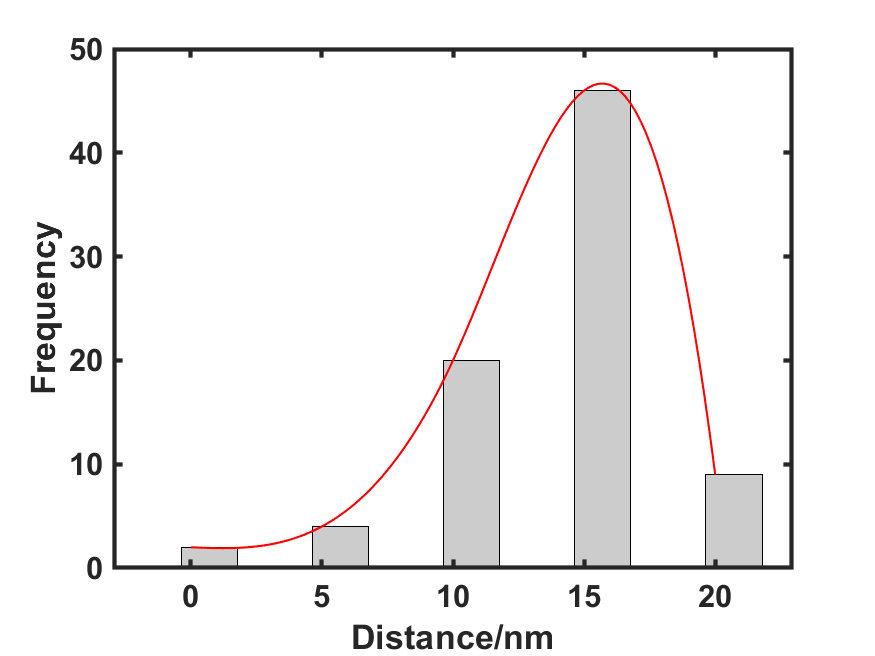

**Figure S2-10.** Statistic distributions of the distance between the assembled SDEM(SOX-HRP) enzyme pairs. The average distance for the two enzymes linked with DNA double strand was ~12 nm.

**Section S3: The Optimization of SDEM Cascade Activity.**

Free enzymes: Free enzymes indicate the freely diffusing enzyme pairs (GOx, HRP), which weren’t modified with DNA. SSDMEs(GOx, HRP): SSDMEs(GOx, HRP) were the freely diffusing enzyme pairs (GOx-L1, HRP-L2), which were modified with DNA. SDEM(GOx-HRP): SDEM(GOx-HRP) were the SSDMEs(GOx, HRP) pairs linked with DNA (L3, A or A’).

The titration curves of three kinds of enzyme pairs revealed the absorbance change of oxidation of TMB. 4 nM free enzymes, SSDMEs(GOx, HRP) and SDEM(GOx-HRP) were diluted to 0.8 nM for activity assays. The activity assays were performed on a SpectraMax iD5 96 well plate reader (Molecular Devices, USA). Enzyme cascade activity was measured in 4 mM glucose and 0.064 g/L TMB substrate (pH 7.4). Enzyme cascade activity was evaluated by the absorbance of oxidation of TMB which was monitored by the change in absorbance at 370 nm. The titration curves of GOx/HRP showed that the SDEM(GOx-HRP) cascade reaction reached equilibrium in 10 min compared to free enzymes and SSDMEs(GOx, HRP) in solution.

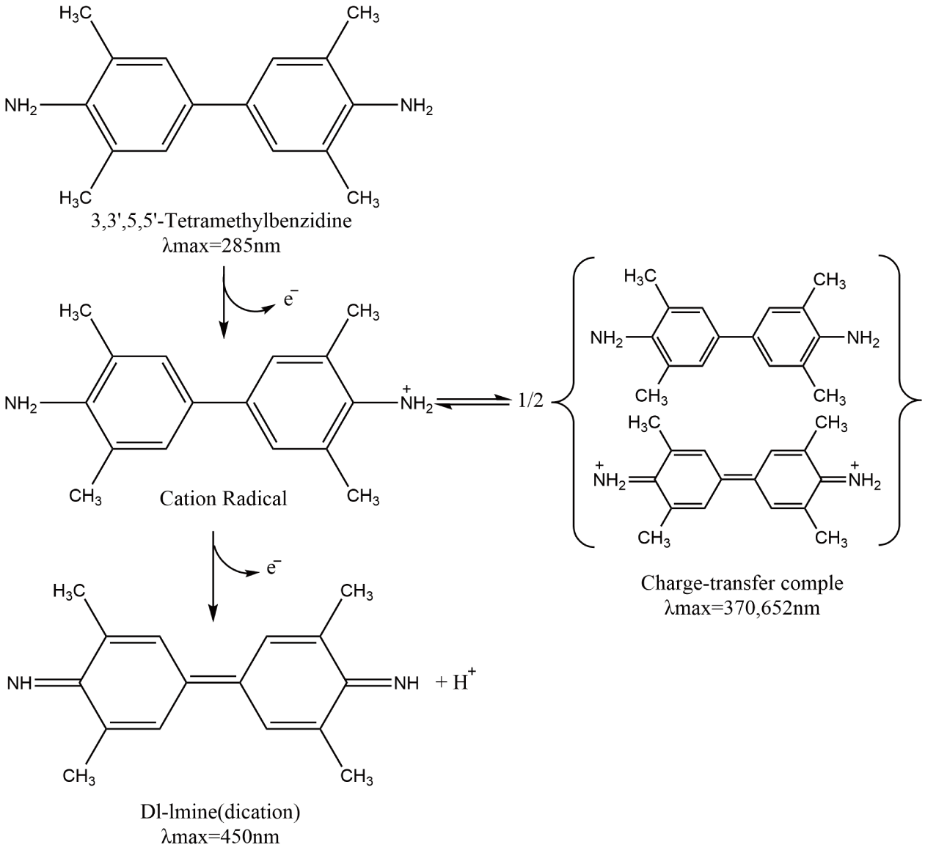

**Figure S3-1.** The charge transfer of oxidation of TMB. In enzyme cascade reaction, TMB was oxidized by hydrogen peroxide and the absorbance of the product was at 370 nm. In the presence of excess hydrogen peroxide, the product at 370 nm was further oxidized and the absorbance at 370 nm was decreased.

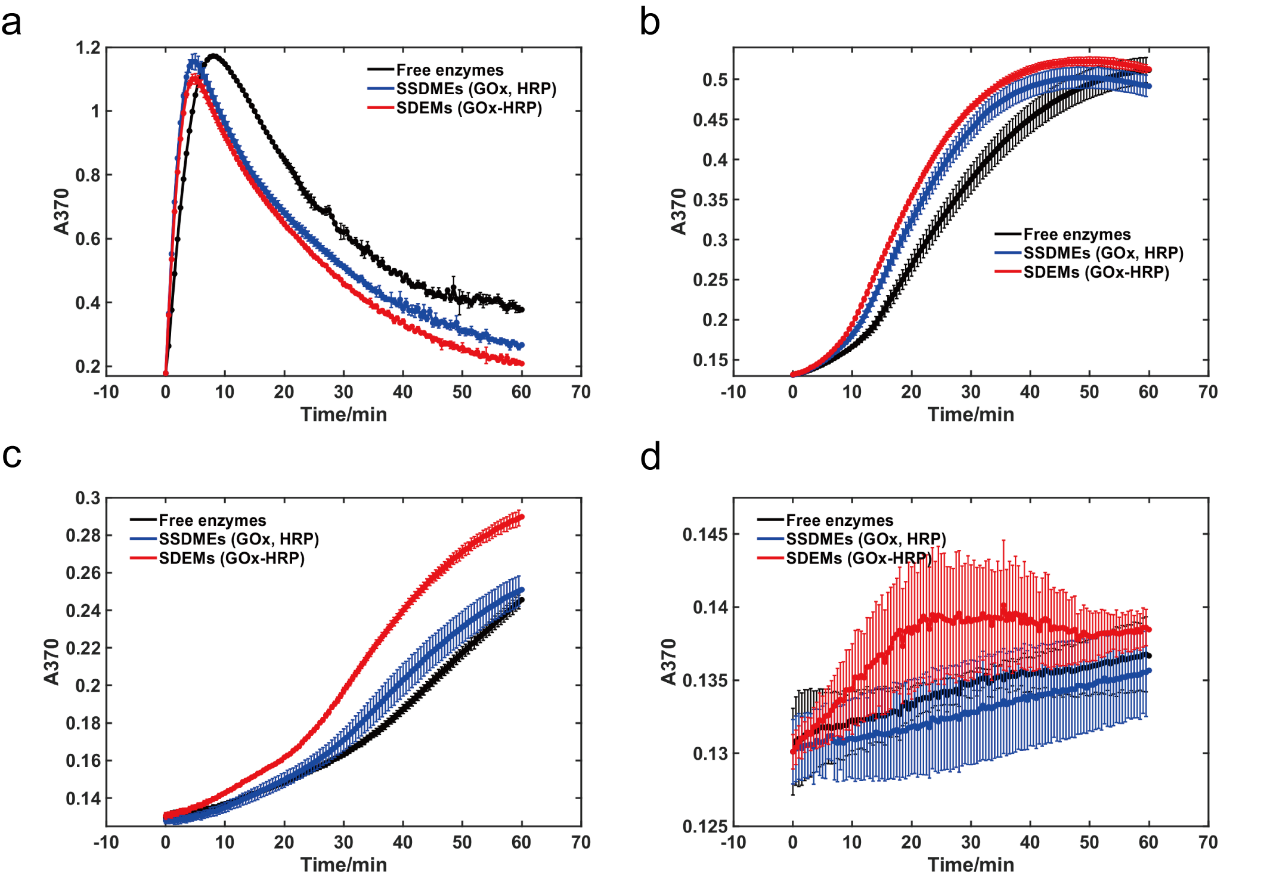

**Figure S3-2.** Enzyme activity for free enzymes, SSDMEs (GOx, HRP) and SDEM(GOx-HRP) at different concentration of the GOx/HRP cascades. (a) 8 nM. (b) 0.8 nM. (c) 0.4 nM. (d) 0.08 nM. At room temperature.

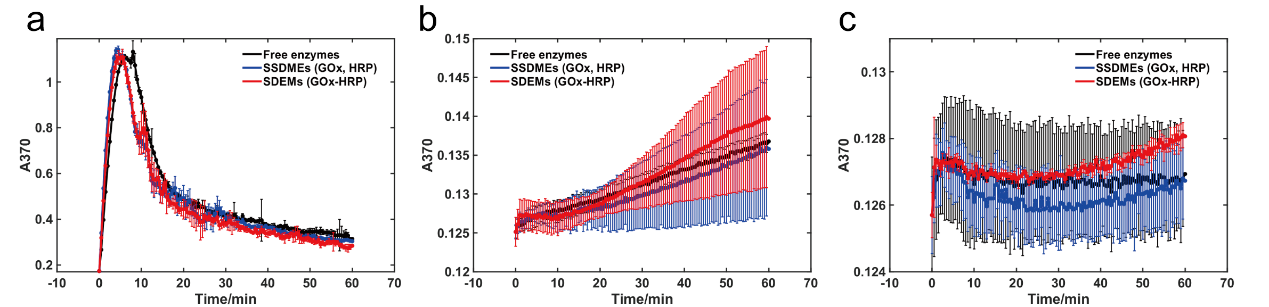

**Figure S3-3.** Enzyme activity for free enzymes, SSDMEs(GOx, HRP) and SDEM(GOx-HRP) at different concentration of the GOx/HRP cascades. (a) 8 nM. (b) 0.4 nM. (c) 0.08 nM. At 37 ℃.

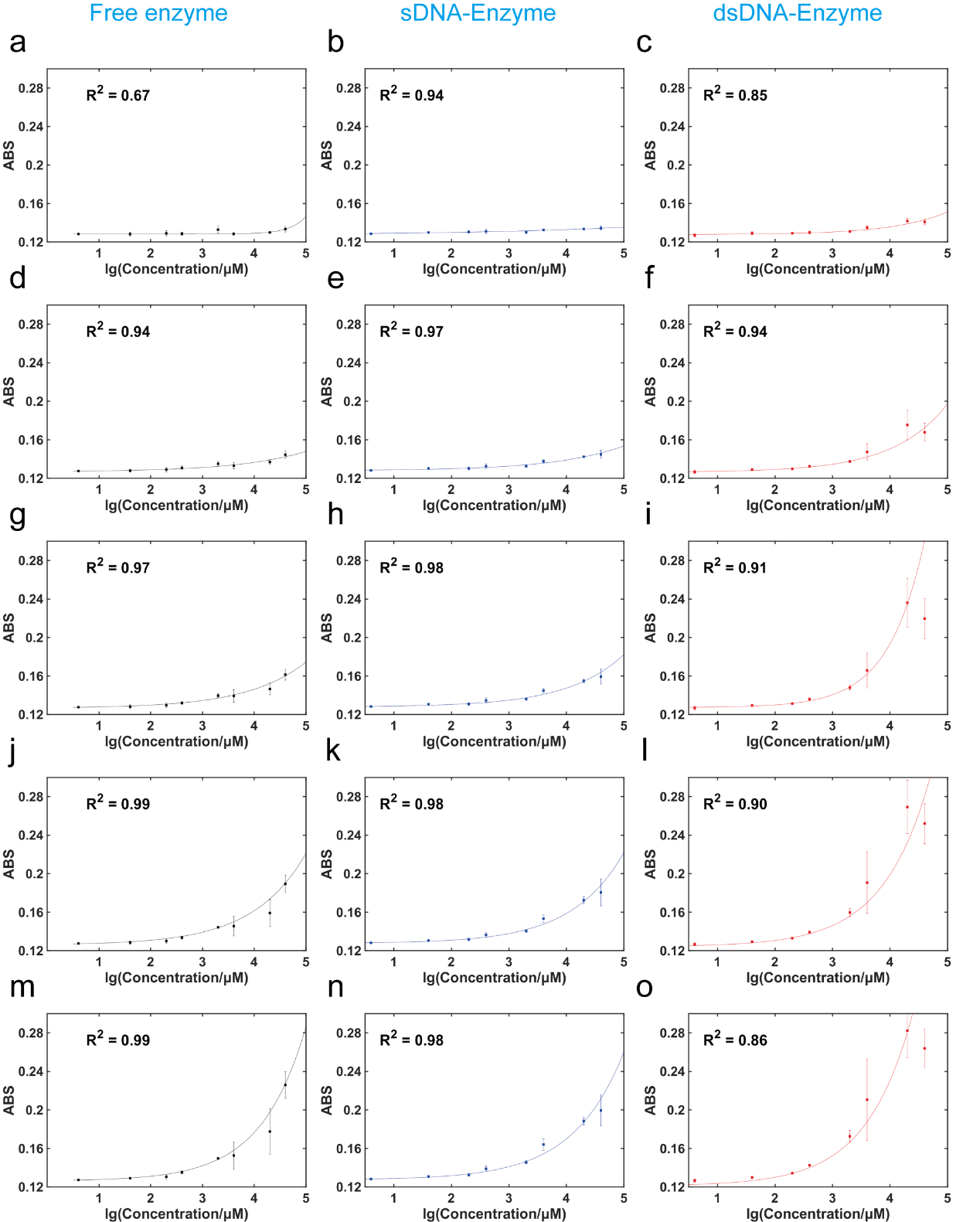

**Figure S3-4.** The titration curve of glucose detection for free enzymes, SSDMEs(GOx, HRP) and SDEM(GOx-HRP) in solutions over 50 min. (a, b, c) 10 min (d, e, f) 20 min (g, h, i) 30 min (j, k, l) 40 min (m, n, o) 50 min. (a, d, g, j, m) Free enzymes (fit line: black color). (b, e, h, k, n) SSDMEs(GOx, HRP) (fit line: blue color). (c, f, i, l, o) SDEM(GOx-HRP) (fit line: red color). Enhancement activity of the SDEM(GOx-HRP) compared to free enzymes and SSDMEs(GOx, HRP) in solution.

**Section S4: Nonspecific DNA Interactions.**

Nonspecific DNA interactions were evaluated in the presence of nonspecific DNA strand G and SDEM (0.97 nM GOx-HRP) in switch regulation. The concentration of nonspecific DNA was equal to the fuel DNA strand F in switch regulation. The ABS showed that nonspecific DNA had no effect on the SDEM(GOx-HRP) cascade reaction as shown in Figure S4-1.

Nonspecific DNA interactions were also evaluated in the SDEM(GOx-HRP) at ultrahigh concentration of nonspecific DNA. 100-fold Excess of nonspecific DNA had an insignificant effect on the SDEM(GOx-HRP) cascade reaction. 10000-fold Excess of nonspecific DNA had an obvious effect on the SDEM(GOx-HRP) cascade reaction as shown in Figure S4-2. At ultrahigh concentration of nonspecific DNA, recently studies on the role of DNA nanostructures in the catalytic properties showed that the electrostatic environment near the encaged enzyme can have an effect on the enzyme activity^7^.

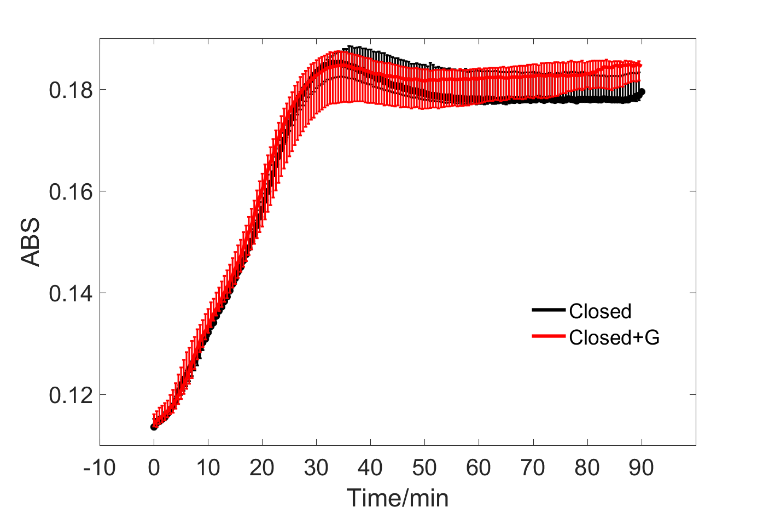

**Figure S4-1.** Nonspecific DNA interactions were performed in the SDEM(GOx-HRP) cascade reaction. Observation of the ABS of SDEM(GOx-HRP) cascade in absence G (black line, 1.0 nM Closed state) and in the presence of 3.6 nM nonspecific DNA strand G (blue line, Closed + G, 3.6-fold excess). Nonspecific DNA had no effect on the SDEM(GOx-HRP) cascade reaction at this concentration.

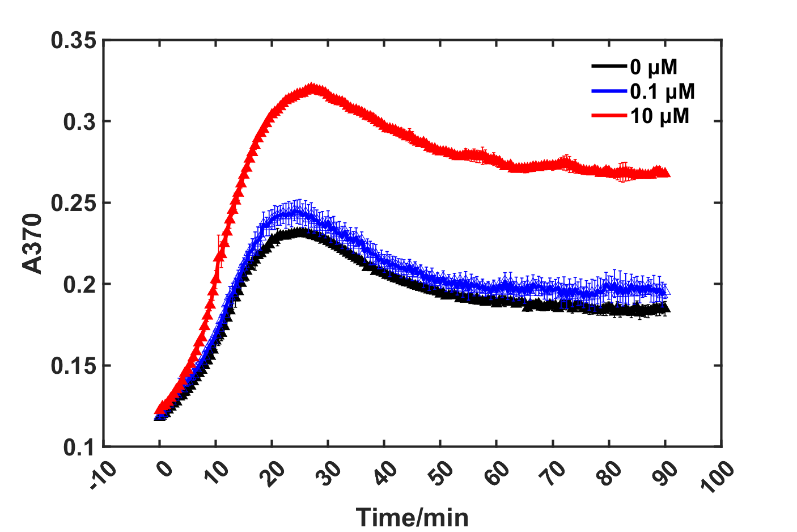

**Figure S4-2.** High concentration of nonspecific DNA interactions was performed in the SDEM(GOx-HRP) cascade reaction. Observation of the ABS of SDEM(GOx-HRP) cascade in absence G (black line, 1.0 nM SDEM(GOx-HRP)), in the presence of 100 nM nonspecific DNA strand G (blue line, 100-fold excess) and 10 μM G (red line, 10000-fold excess). High concentration nonspecific DNA strands influenced the enzymes activities of the SDEM(GOx-HRP).

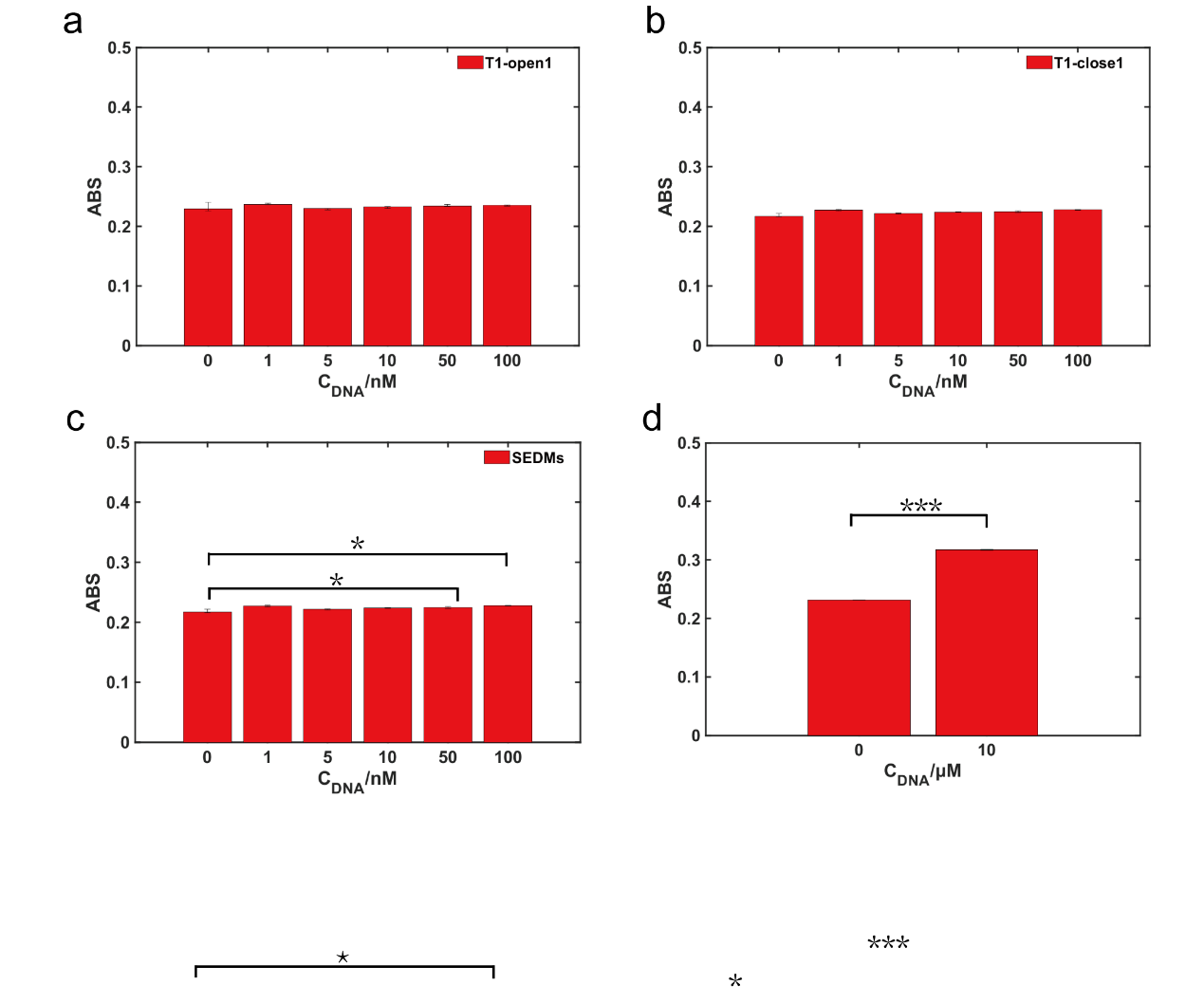

**Figure S4-3.** Elevated the influence of nonspecific DNA in T1-open1 (SDEM, 1 nM GOx-HRP), T1-close1 (SDEM, 1 nM GOx-HRP) and SDEM (1 nM GOx-HRP) at different concentration of GO compared with the control (0 nM GO). (a) T1-open1. (b) T1-close1. (c)(d) SDEM(GOx-HRP).
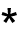
p<0.05;
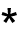

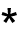

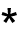
p<0.001.

**Section S5: Verifying the Enhancement SDEM Cascade Reactions for Sarcosine Detection.**

Free enzymes: Free enzymes were the freely diffusing enzyme pairs (SOX, HRP), which weren’t modified with DNA. SDEM(SOX-HRP): SDEM(SOX-HRP) was the SSDMEs(SOX, HRP) pairs linked with DNA (L3).

1 μM SOX/HRP were diluted to 200 nM for activity assays and enzyme cascade activity was measured in 0.064 g/L TMB substrate (pH 7.4) at room temperature for 30 min.

Remarkable increase was observed in ABS from 8 μM to 40 μM sarcosine comparing with blank group (0 μM sarcosine) as shown in Figure S5-1. In this concentration range of sarcosine, free enzymes and SDEM(SOX-HRP) cascade reaction were sensitive.

At the different concentration of enzymes, the SOX/HRP enzyme activity assays of free enzymes and SDEM(SOX-HRP) were performed. SDEM(SOX-HRP) pairs enhanced the activity of enzyme cascade compared to free enzymes in solution as shown in Figure S5-2 which contributed to the sensitivity of SDEM(SOX-HRP).

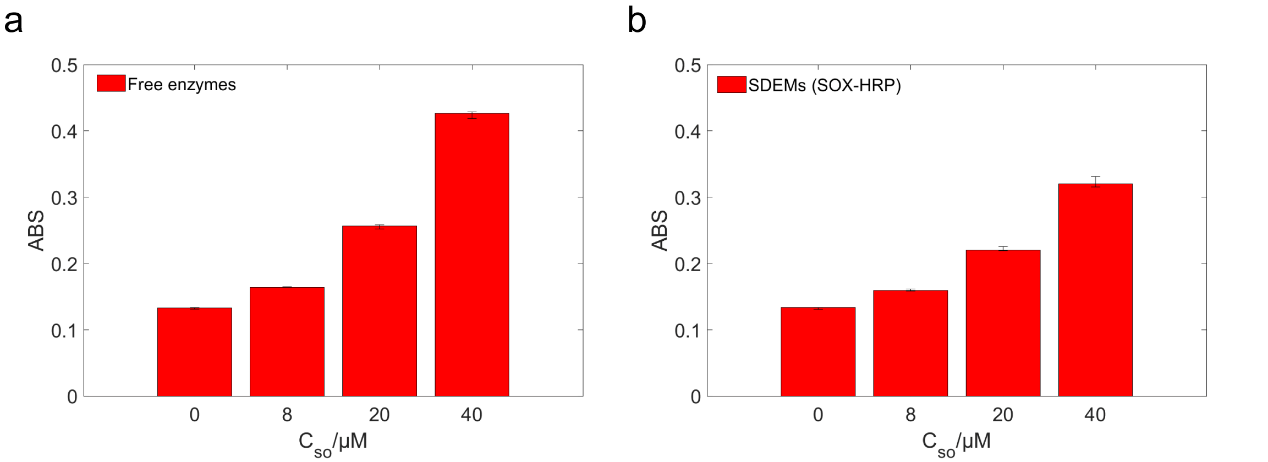

**Figure S5-1.** The enzymes activities of free enzymes and SDEM(SOX-HRP). (a) The enzymes activities detected via TMB’s Abs of 200 nM free enzymes in different concentration SO. (b) The enzymes activities detected via TMB’s Abs of 200 nM SDEM(SOX-HRP) in different concentration SO.

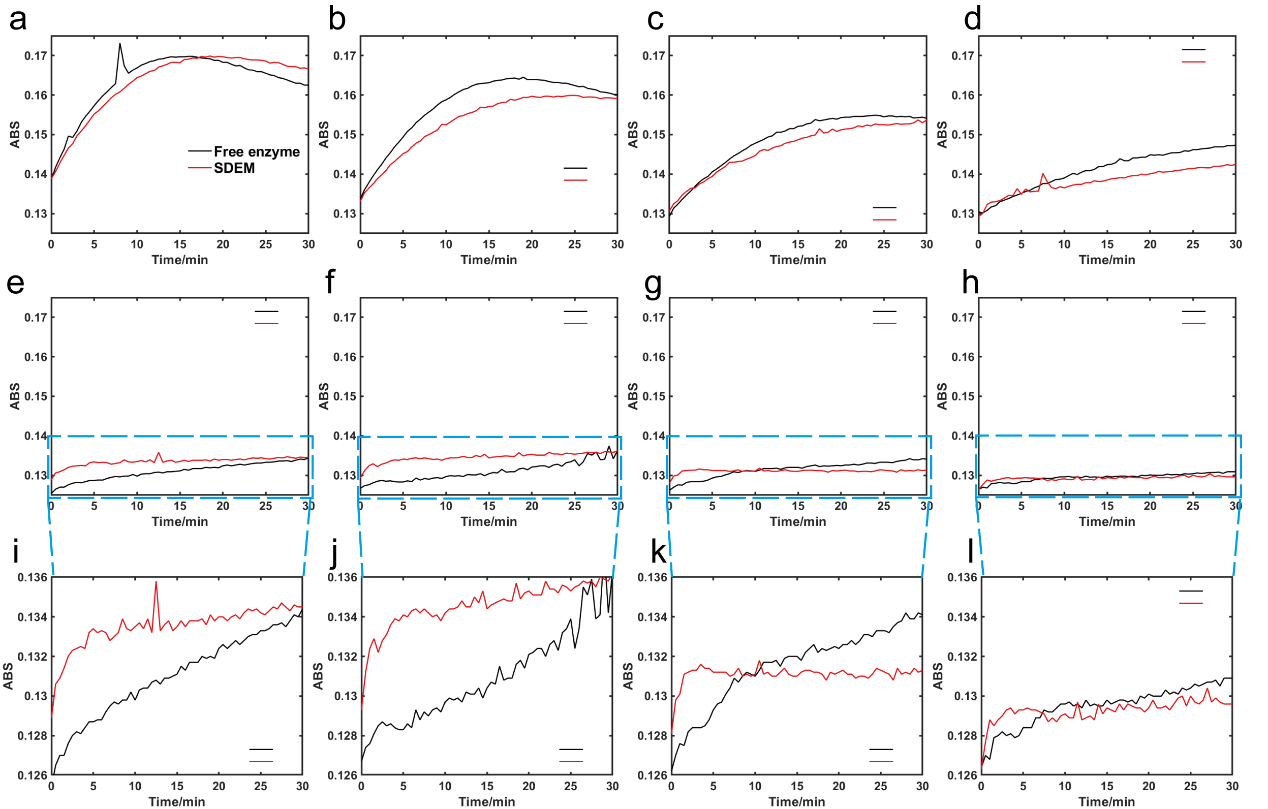

**Figure S5-2.** The SOX/HRP enzyme activity of free enzymes and SDEM(SOX-HRP) under the different concentration of enzymes. (a) 160 nM. (b) 120 nM. (c) 80 nM. (d) 40 nM. (e) 20 nM. (f) 16 nM. (g) 12 nM. (h) 8 nM. (i) (j) (k) (l) were the higher resolution images of (e) (f) (g) (h), respectively. (i) (j) (k) (l) showed enhanced activity of the SDEM(SOX-HRP) compared to free enzymes in solution.

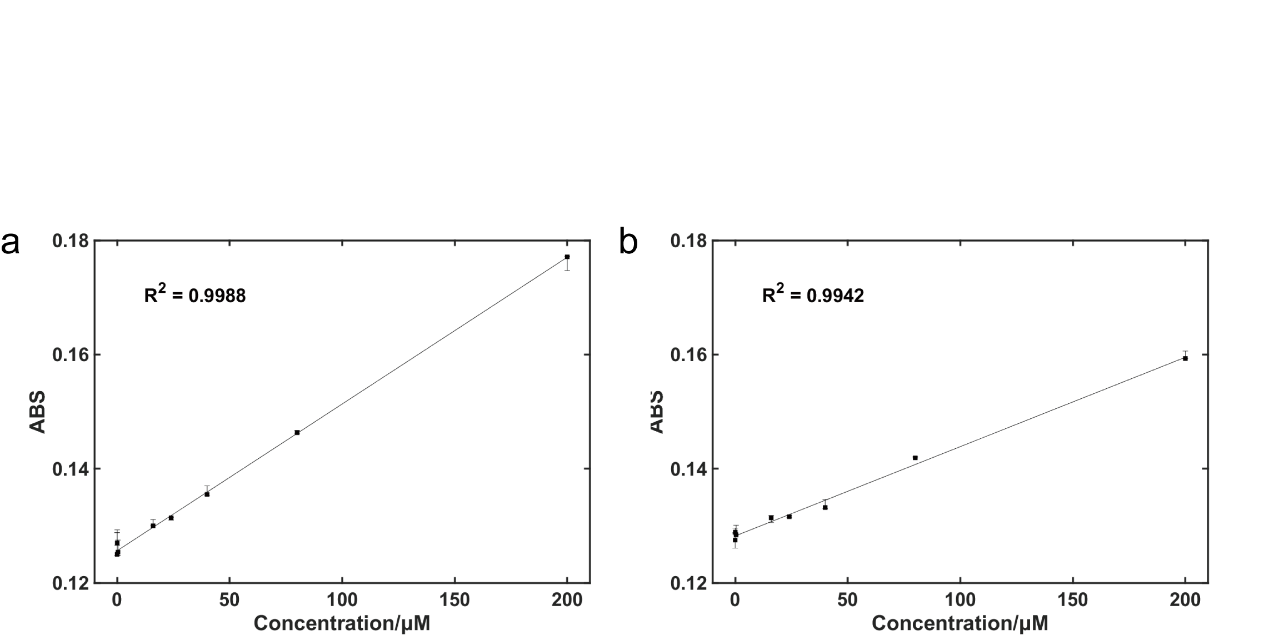

**Figure S5-3.** Comparison of observed and the titration curve of sarcosine detection for free enzymes and SDEM(SOX-HRP) in solutions with 20 nM SOX/HRP. (a) Free enzymes. (b) SDEM(SOX-HRP). SDEM(SOX-HRP) cascade reaction had high sensitivity (16 μM) compared to free enzymes (40 μM).

**Section S6: The Programable SDEMs.**

**The design of the programable SDEMs**. Programable switch was designed from the principle for highly specific switch regulation based on a DNA-fuelled molecular machine^8^. The programable SDEMs were closed or opened with excessive additions of D/F or the corresponding complementary strand of D/F respectively. Two alternatives were designed in this work and were different in the stem energy.

**The programable SDEMs with -6 kcal mol^-1^ stem energy.** In the first scheme, the programable SDEMs were prepared by mixing stoichiometric quantities of the three strands, A1, B and C, in TM buffer at 95 ℃ for 10 min, then cooled to 4 °C over 30 s. The programable SDEM was opened with fuel strands D (1.2-fold excess for A1+B+C, shown in Figure S6-1a) and the ΔΔG associated with the hybridization of a complementary base pair was -24 kcal mol^-1^ at 25 ℃ (shown in Figure S6-1b(1)). The open state returned to the closed state with fuel strands E (1.2-fold excess for D) or D* (1.2-fold excess for D) to form a double-stranded waste product DE or DD* and the ΔΔG were -10 kcal mol^-1^ (shown in Figure S6-1b(2)) and -20 kcal mol^-1^ (shown in Figure S6-1b(3)) at 25 ℃, respectively.

**Characterization of the yield of the programable SDEMs with -6 kcal mol^-1^ stem energy.** PAGE showed the construction of DNA molecular machine and the regulation of programable switch as shown in Figure S6-1c. Lane 2 - 4 were the construction of SDEMs (closed state). From lane 4 to 5, the strand D hybridized to the stem of the closed state and the closed state was opened. From the gray value of the lane 5, the closed state of SDEMs was opened with fuel strands D in yields of 50%. From the gray value of the lane 6, the open state returned to the closed state with fuel strands E/D* in yields of 100%. Since the design of SDEMs from the closed state to open state just in yields of 50%, the design scheme could be further optimized.

**The programable SDEMs with -2 kcal mol^-1^ stem energy.** To increase the yield of the programable switch’s opening, a second assembled programable switch was designed with the lower stem energy (-2 kcal mol^-1^). The second assembled programable switch was prepared by mixing stoichiometric quantities of the three strands, A2, B and C, in TM buffer at 95 ℃ for 10 min, then cooled to 4 °C over 30 s. The second assembled programable switch were closed and opened with fuel strands as shown in Figure S6-2a. The second assembled programable switch were opened with fuel strands F (1.2-fold excess for the second assembled programable switch) and the ΔΔG associated with the hybridization of a complementary base pair was -25 kcal mol^-1^ at 25 ℃ as shown in Figure S6-2b(1). The open state returned to the closed state with fuel strands F* (1.2-fold excess for F) to form a double-stranded waste product FF* and the ΔΔG were -16 kcal mol^-1^ as shown in Figure S6-2b(2) at 25 ℃.

**Characterization of the yield of the programable SDEMs with -2 kcal mol^-1^ stem energy.** The second assembled programable switch were opened and the open state returned to the closed state in yields of 100% and 100% (shown in Figure S6-2c). The scheme of the second assembled programable switch was applied to characterize the state of the programable switch and the study of the enzyme cascade reaction.

**Fluorescence characterization of the programable SDEMs.** Dye quenching was used to characterize the state of the second assembled programable switch in Figure S6-3. The fluorescence intensity increased by a factor of seven when the programable SDEMs were opened, and went back to approximately the same level when the open state returned to the closed state.

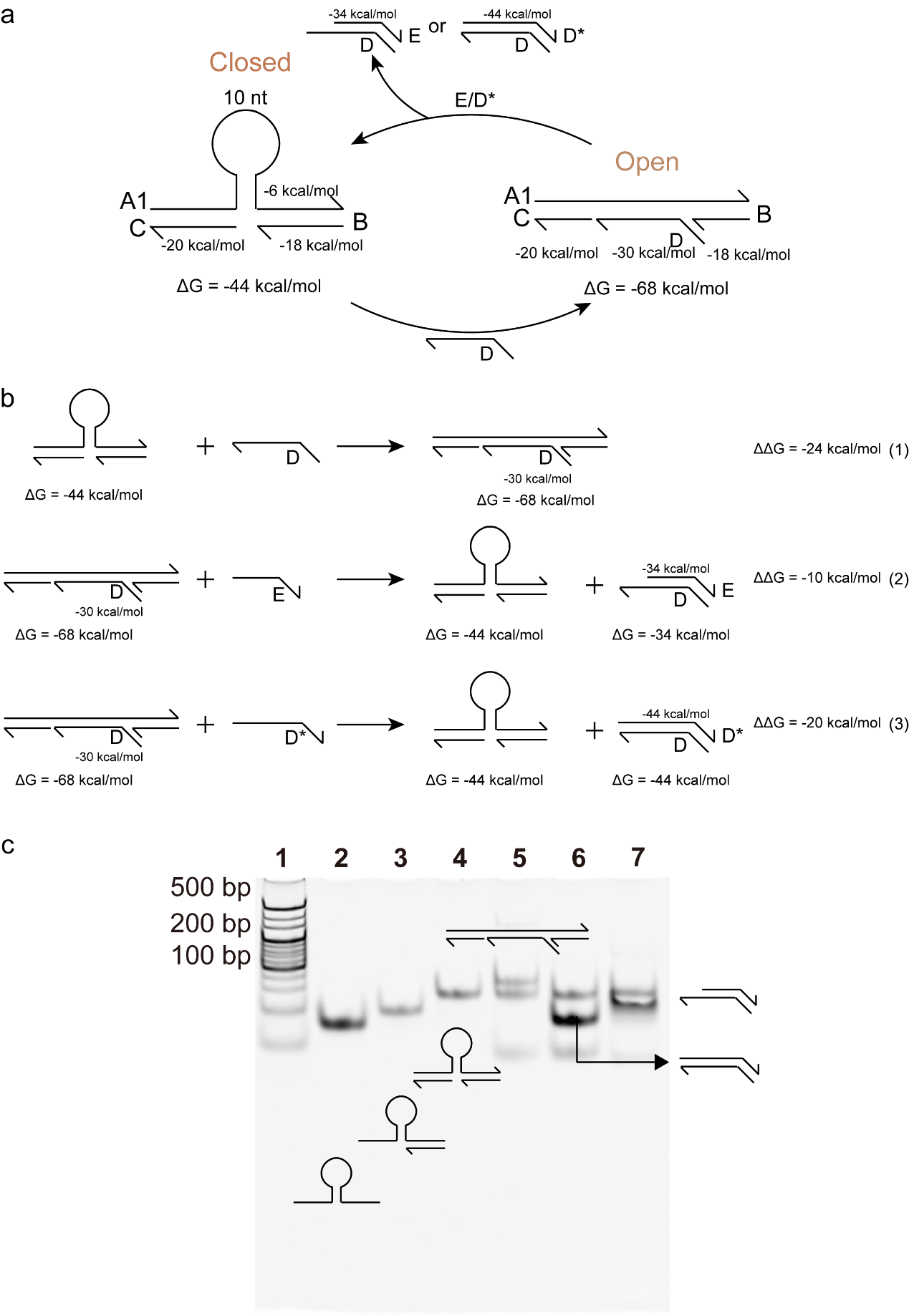

**Figure S6-1.** Programable switch design with -6 kcal mol^-1^ stem energy. (a) Construction and operation of the programable SDEMs. (b) The ΔΔG associated with the operation of the programable SDEMs. (c) Analysis of the programable SDEMs formation by PAGE. Lane 1: ladder; lane 2: A1; lane 3: A1+B; lane 4: A1+B+C; lane 5: A1+B+C+D; lane 6: (A1+B+C+D) +E; lane 7: (A1+B+C+D) +D*.

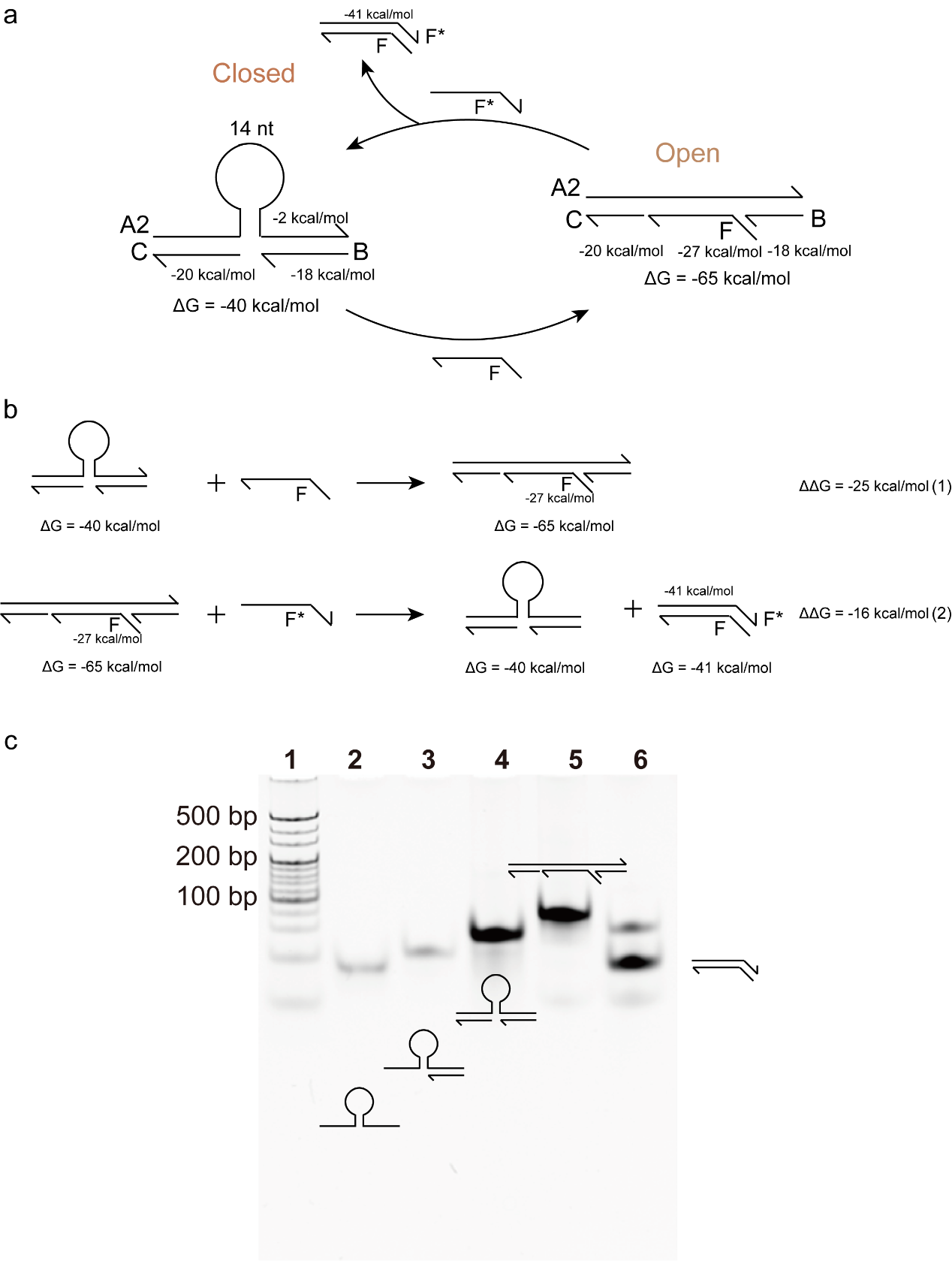

**Figure S6-2.** Programable switch design with -2 kcal mol^-1^ stem energy. (a) Construction and operation of the second assembled programable switch. (b) The ΔΔG associated with the operation of the second assembled programable switch. (c) Analysis of the second assembled programable switch formation by PAGE. Lane 1: ladder; lane 2: A2; lane 3: A2+B; lane4: A2+B+C; lane 5: A2+B+C+F; lane 6: (A2+B+C+F) +F*. This is the detailed description of the gel in Fig. 4f.

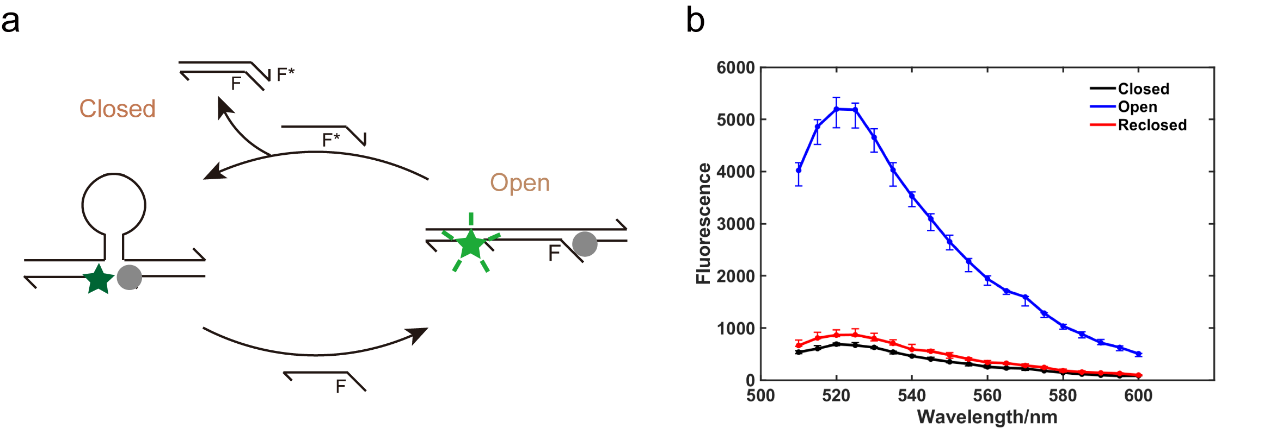

**Figure S6-3.** Fluorescence characterization of the programable SDEMs. (a) Schematic illustration of the change of fluorescence intensity. (b) The DNA strand B was labelled at the 3’ ends with dyes Black Hole Quencher-1 (BHQ1) and the DNA strand C was labelled at the 5’ ends with dyes FAM. When FAM was excited by the 480 nm emission, it fluoresces with a peak emission wavelength of 520 nm. This emission was quenched by fluoresce resonant energy transfer (FRET) from FAM to BHQ1 (“closed programable switch”) and the efficiency of FRET decreases as the distance between the dyes increases (“opened programable switch”). The fluorescence intensity showed that by adding excessive closing and opening strands in sequence, and the programable switch can be closed and opened.

**Section S7: The SDEM Cascade Reactions and Networks Catalyzed by DNA**

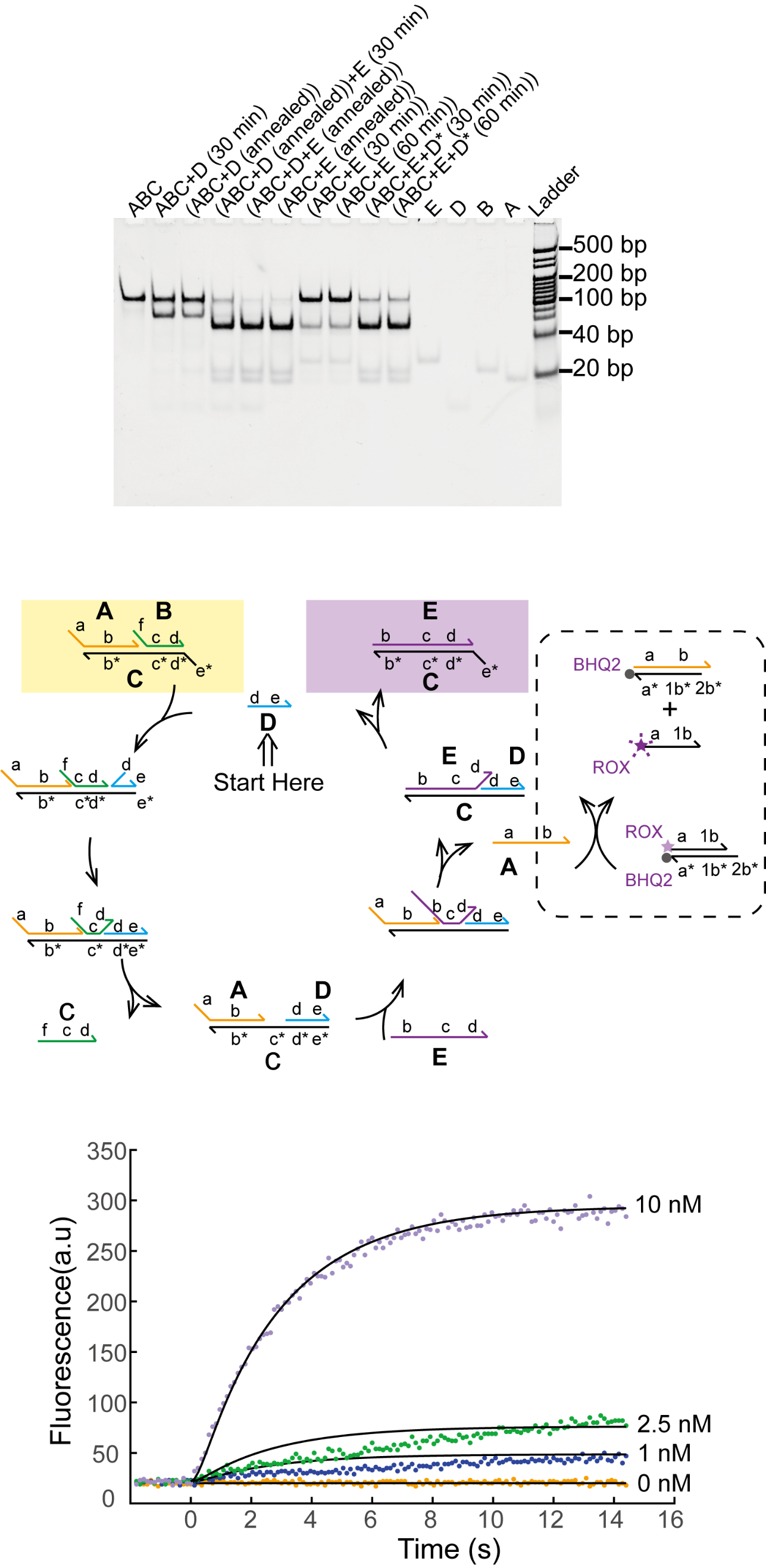

**Figure S7-1.** Gel results of the cascade reactions and networks catalyzed by DNA. All experiments were performed at 25°C in tris-acetate (TE with 12.5 mM MgCl_2_), unless otherwise noted. [A] = [B] = [C] = [D] = 10 nM, [D*] = [E]= 200 nM. All annealing processes were performed at 95 °C to 20 °C at a constant rate over the course of 90 minutes. Conditions: 12% PAGE with a constant voltage of 180 V for 60 mins.

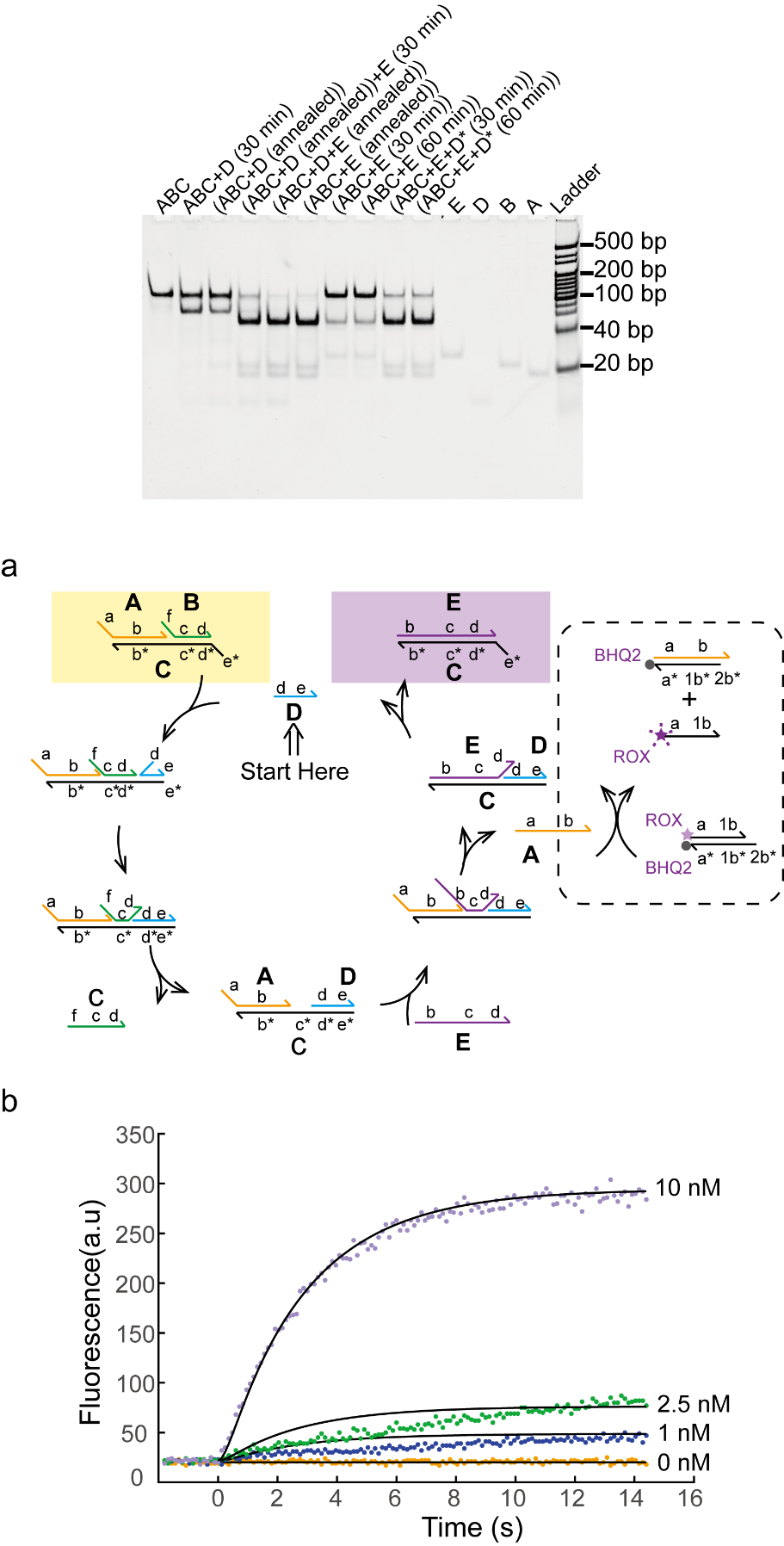

**Figure S7-2.** Demonstration of catalysis. (a) Schematic of the catalytic pathway without enzymes. Schematic of output (the DNA strand a-b) reporter using 6-carboxy-X-rhodamine (ROX) and Black Hole Quencher-2 (BHQ2). (b) Fluorescent results of the cascade reactions and networks catalyzed by DNA. Different concentration of the catalyst strand D was introduced into the system. The orange trace, blue trace, green trace and purple trace were investigated with 0 nM, 1 nM, 2.5 nM and 10 nM catalyst strand. [ABC] = 10 nM, [(a-1b) - (2b*-1b*-a*)] = 30 nM, [E] = 20 nM. Dotted lines showed the experiment data. The solid line showed simulation data.

The assembly yields of SDEM(GOx-HRP) and SDEM(SOX-HRP) were evaluated with the SDEM cascade reactions catalyzed by catalytic DNA strand. GOx-HRP was a two-enzymes (GOx and HRP), a linked double-stranded DNA. SOX-HRP was a two-enzymes, SOX and HRP, a linked double-stranded DNA. SSDMEs(GOx, HRP) and SSDMEs(SOX, HRP) were the free enzymes modified with DNA.

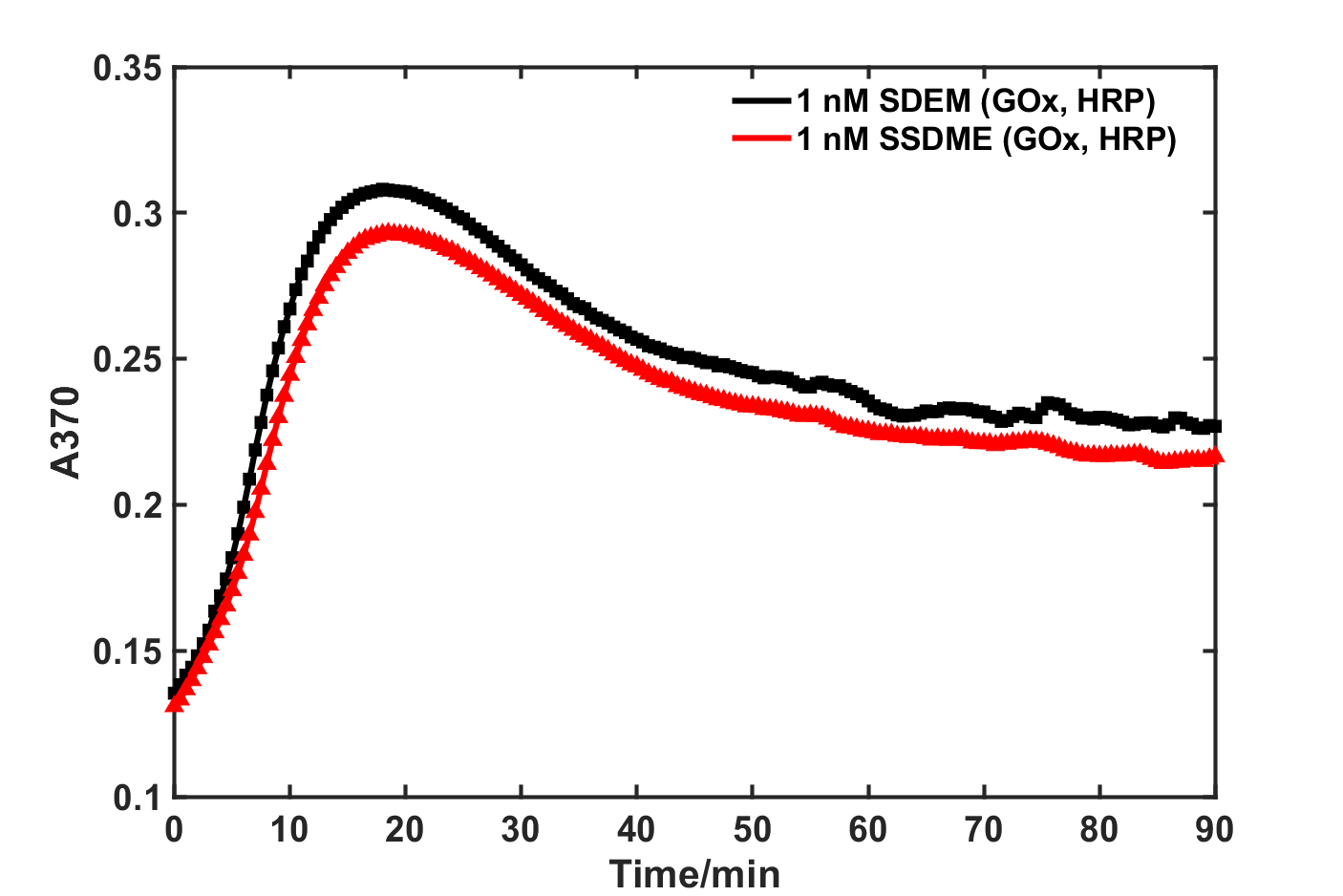

**Figure S7-3.** The titration curves of the 1 nM SDEM(GOx-HRP) and 1 nM SSDMEs(GOx, HRP). The titration curves showed that the enzyme cascade activity of SDEM(GOx, HRP) was higher than the SSDMEs(GOx, HRP).

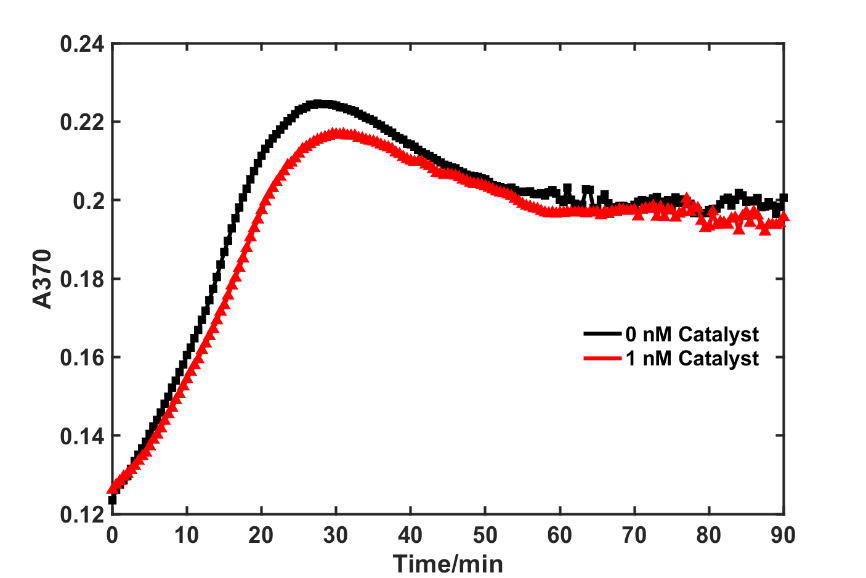

**Figure S7-4.** The titration curves of the 1 nM GOx-HRP with DNA catalyst (the DNA strand D) at the concentration of 0 nM and 1 nM. When the DNA strand D were added into the GOx-HRP, the enzymes GOx were released and moved far away from the enzymes HRP so that the enzyme cascade activity of GOx-HRP was decreased. The assembly yield of SDEM(GOx-HRP) can be approximately estimated by comparing the decrease of the absorbance in Figure S6-1 with in Figure S6-2 using equation in Table S6-1. The assembled SDEM(GOx-HRP) were in yield of 64.10%.

**Table S7-1.** The assembly yield of SDEM(GOx-HRP) at the concentration of 1 nM (n = 3 samples; means).

| **A_0(GOx-HRP)_** | **A_1(GOx+HRP)_** | **ΔA_1_(A_0_-A_1_)** | **A_0_’_(GOx-HRP)_** | **A_2(GOx+HRP)_** | **ΔA_2_(A_0_’- A_2_)** | **Yield(ΔA_2_/ΔA_1_)** |
| --- | --- | --- | --- | --- | --- | --- |
| 0.3261 | 0.3030 | 0.02310 | 0.2031 | 0.1883 | 0.01480 | 64.10% |

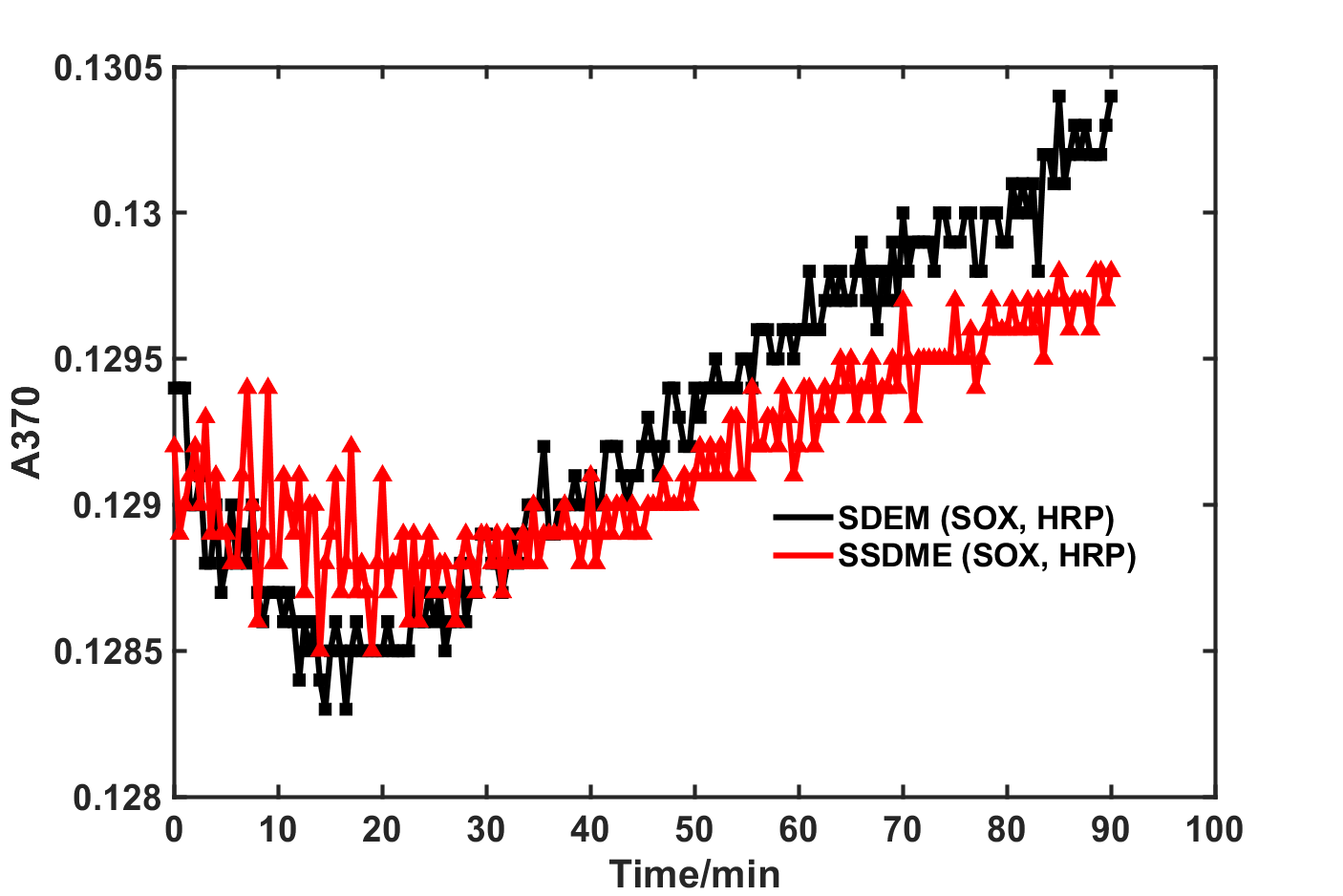

**Figure S7-5.** The titration curves of the 20 nM SDEM(SOX-HRP) and 20 nM SSDMEs(SOX, HRP). The titration curves indicated that the enzyme cascade activity of SDEM(SOX-HRP) was higher than the SSDMEs(SOX, HRP).

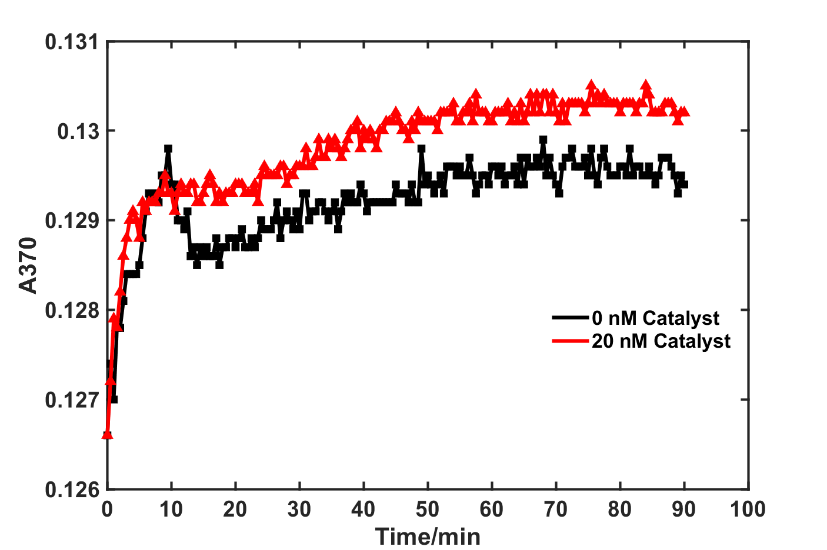

**Figure S7-6.** The titration curves of the 20 nM SSDMEs(SOX,HRP) with catalyst DNA (the DNA strand D) at the concentration of 0 nM and 20 nM . When the DNA strand D were added into the SSDMEs(SOX,HRP), the SOX were assembled with the HRP so that the enzyme cascade activity of SSDMEs(SOX,HRP) was increased. The assembly yield of SDEM(SOX-HRP) can be approximately estimated by comparing the increase of the absorbance in Figure S6-3 with in Figure S6-4 using equation in Table S6-2. The assembled SDEM(SOX-HRP) were in yield of 62.50%.

**Table S7-2.** The yield of the assembled SDEM(SOX-HRP) at the concentration of 20 nM (n = 3 samples; means).

| **A_0(SOX-HRP)_** | **A_1(SOX+HRP)_** | **ΔA_1_(A_0_-A_1_)** | **A_0_’_(SOX-HRP)_** | **A_2(SOX+HRP)_** | **ΔA_2_(A_0_’- A_2_)** | **Yield(ΔA_2_/ΔA_1_)** |
| --- | --- | --- | --- | --- | --- | --- |
| 0.1304 | 0.1296 | 0.0008 | 0.1300 | 0.1295 | 0.0005 | 62.5% |

**Section S8: DNA sequence.**

Table S8-1. Oligos used in the SDEM cascade reactions compared with free enzymes and SSDMEs.

| **Oligos name** | **Sequence (5’ to 3’)** |
| --- | --- |
| L1-SH | SH-TTTTTTTTTTTTTTTTTTTTTT |
| L2-SH | TTGGTGGTGGTGGTGGTGGT-SH |
| L3 | ACCACCACCACCACCACCAAAAAAAAAAAAAAAAAAAAAAAA |
| L1 | TTTTTTTTTTTTTTTTTTTTTT |
| L2 | TTGGTGGTGGTGGTGGTGGT |

Table S8-2. Oligos used in the programable SDEM.

| **Oligos name** | **Sequence (5’ to 3’)** |
| --- | --- |
| A1 | CAGTCCACAAGCTATCGAGTCGATCAACCATCCGACTCACAAGAGCAATCAATA |
| 4-2-3-2*-1 (A2) | CAGTCCACAAGCTATC-GAGT-ATATCAACCATCCG-ACTC-ACAAGAGCAATCAATA |
| 1*-SH | SH-TATTGATTGCTCTTGT |
| 4*-SH | GATAGCTTGTGGACTG-SH |
| D | AGTACTTACTGGAGTCGGATGGTTGATCGACTC |
| D* | GAGTCGATCAACCATCCGACTCCAGTAAGTACT |
| E | TCAACCATCCGACTCCAGTAAGTACT |
| 1* (B) | TATTGATTGCTCTTGT |
| 4* (C) | GATAGCTTGTGGACTG |
| 4*-FAM | FAM-GATAGCTTGTGGACTG |
| 1*-BHQ1 | TATTGATTGCTCTTGT-BHQ1 |
| 6-2*-3-2-1 | CAGTCCACAAGCTATC-GAGT-ATATCAACCATCCG-ACTC-TTGGTGGTGGTGGTGG |
| 4-2*-3-2-7 | AGTGCTGCCATCTACT-GAGT-ATATCAACCATCCG-ACTC-ACAAGAGCAATCAATA |
| 9-2*-3-2-8 | GCTACCTCCTAGTGAC-GAGT-ATATCAACCATCCG-ACTCTCTCACGTCTGCTCTC |
| 4-2*-3-2-10 | CGAGACCAGCCTGGCC-GAGT-ATATCAACCATCCG-ACTC-ACAAGAGCAATCAATA |
| 6*-7* | GATAGCTTGTGGACTG-TATTGATTGCTCTTGT |
| 6*-8* | GATAGCTTGTGGACTG-GAGAGCAGACGTGAGA |
| 9*-10* | GTCACTAGGAGGTAGC-TATTGATTGCTCTTGT |
| 5-2*-3*-2 | AGTACTTACTG-GAGT-CGGATGGTTGATAT-ACTC |
| 2*-3-2-5* | GAGT-ATATCAACCATCCG-ACTC-CAGTAAGTACT |

Table S8-3. Oligos used in the SDEM cascade reactions and networks catalyzed by DNA.

| **Oligos name** | **Sequence (5’ to 3’)** |
| --- | --- |
| (f-c-d)-SH | CCACATACATCATATT-CCCT-CATTCAATACCCTACG-SH |
| (b-c-d)-SH | CCTACGTCTCCAACTAACTTACGG-CCCT-CATTCAATACCCTACG-SH |
| (e*-d*-c*-b*)-SH | TGGAGA-CGTAGGGTATTGAATG-AGGG-CCGTAAGTTAGTTGGAGACGTAGG-SH |
| d-e | CATTCAATACCCTACG-TCTCCA |
| a-b | CTTTCCTACA-CCTACGTCTCCAACTAACTTACGG |
| f-c-d | CCACATACATCATATT-CCCT-CATTCAATACCCTACG |
| b-c-d | CCTACGTCTCCAACTAACTTACGG-CCCT-CATTCAATACCCTACG |
| e*-d*-c*-b* | TGGAGA-CGTAGGGTATTGAATG-AGGG-CCGTAAGTTAGTTGGAGACGTAGG |
| CAOR1-ROX | ROX-CTTTCCTACACCTACG |
| CAOR2-BHQ2 | TGGAGACGTAGGTGTAGGAAAG-BHQ2 |
| CASR1-FAM | FAM-CCACATACATCATATTCCCT |
| CASR2-BHQ1 | TTGAATGAGGGAATATGATGTATGTGG-BHQ1 |
| 1b* | CGTAGG |
| 2b* | TGGAGA |
| 3b* | CCGTAAGTTAGT |

Table S8-4. Nonspecific DNA Interactions.

| **Oligos name** | **Sequence (5’ to 3’)** |
| --- | --- |
| G | GGAGGAGCCAGAGAAAGTAAATATG |

7.	The role of DNA nanostructures in the catalytic properties of an allosterically regulated protease. *Science Advances* (2022).

8.	Yurke, B., Turberfield, A.J., Mills, A.P., Simmel, F.C. & Neumann, J.L. A DNA-fuelled molecular machine made of DNA. *Nature* **406**, 605-608 (2000).
